# Sense codon-misassociated eRF1 elicits ribosome stalling and induction of quality control

**DOI:** 10.1101/2024.09.01.610654

**Authors:** Peixun Han, Mari Mito, Akira Yamashita, Takuhiro Ito, Shintaro Iwasaki

## Abstract

The rescue pathway of stalled ribosomes and ribosome-associated quality control (RQC) serves as a surveillance system for aberrant translation that senses ribosome collisions. Although the molecular mechanism has been extensively studied, the endogenous contexts of ribosome collision rescued by the system are poorly understood. Here, beginning with a study of the codon specificity of the eukaryotic translation termination factor eRF1, we show that transient binding of eRF1 to the UUA sense codon leads to ribosome collision and provides a constitutive source of rescue substrates in humans. eRF1-selective Monosome-Seq and Disome-Seq revealed that eRF1 was recruited not only to stop codons but also to near-cognate sense codons, including the UUA codon. The eRF1 on UUA codons delays translation elongation but does not trigger the termination reaction. Remarkably, Disome-Seq with the depletion of ASCC3 and 4EHP, key factors for the ribosome rescue and translation initiation repression, showed that ribosomes stalled at UUA codons consist of a subpopulation suppressed by those factors. Failure of ribosome collision repression by ASCC3-4EHP triggers stress response, such as expression of the stress-induced transcription factor ATF3. This study highlights the impact of sense codon misrecognition by the termination factor on translation homeostasis in human cells.

## Introduction

mRNA translation is subject to surveillance systems to maintain cellular homeostasis ^1–4^. Extensive efforts over the last decade have revealed the surveillance pathway associated with ribosome collision ^1–4^. In typical growing human cells, ribosomes are sparsely located every 200–400 nt in mRNAs ^5–10^. However, once the ribosome encounters obstacles to its traversal along open reading frames (ORFs), it may pause in the middle of nascent chain elongation. Consequently, trailing ribosomes queue up to the leading ribosomes to generate colliding di-ribosomes or disomes. By recruiting specific factors, disomes function as platforms to induce downstream signaling ^1–4^.

Ribosome rescue and concomitant ribosome-associated quality control (RQC) are one such mechanism. In this pathway, a disome recruits the E3 ligase ZNF598 (Hel2 in yeast), which recognizes the 40S‒40S interface between two collided ribosomes ^11–18^ and ubiquitinates ribosomal proteins (eS10 and uS10 in mammals; uS3 and uS10 in yeast) ^13–15,19,20^. This ubiquitin modification allows the helicase ASCC3 (Slh1 in yeast) to split the leading ribosome in an ATP-dependent manner, permitting the trailing ribosome to resume elongation ^13,17,18,21–23^. Simultaneously, colliding ribosomes trigger the repression of translation initiation through the scaffold protein GIGYF2 and the initiation repression factor 4EHP (also known as eIF4E2) ^24–28^. The temporal reduction in ribosome loading onto the transcript could prevent further ribosome collision. The nascent peptide located on the split 60S is ubiquitinated by the E3 ligase Listerin (Ltn1 in yeast) and NEMF (Rqc2 in yeast) and degraded by proteasome ^11,29–31^, with the aid of Ala-Thr tailing at the C-terminus of the nascent peptide (CAT-tailing), which pushes out lysines in the nascent chain embedded in ribosome exit tunnel for the access to Listerin ^31–33^. CAT tails may function as a degron via a Ltn1-independent mechanism ^34^.

The discovery of this mechanism was based on reporter systems, which induce ribosome stalling ^11,35^. The endogenous target sites rescued by the system were subsequently identified, including *SDD1* mRNA in yeasts ^17^, *XBP1u* mRNA in humans ^36^, and *C-I30* mRNA in flies ^37^. More broadly, poly-Lys motifs are natural substrates for the ribosome rescue system in yeasts ^38^, whose demand increases with age ^39^. Additionally, yeast mRNA surveillance is responsible for clearing chemically damaged mRNAs ^40^. However, our knowledge of the endogenous mRNAs targeted by the ribosome rescue system still needs to be completed, especially in human cells.

Recent studies have shown that stop codons are generally a disome-rich position ^36,38,41^. Translation termination is mediated by eRF1 in eukaryotes. Unlike bacteria, which have two termination factors (RF1 and RF2), eRF1 is responsible for recognizing all three stop codon species (UAA, UAG, and UGA) ^42^. During termination, eRF1 is delivered to the stop codon at the A site, forming a complex with the GTPase eRF3 ^43–45^. eRF3 hydrolyzes GTP, dissociates from eRF1 ^46–48^, and induces conformational change of eRF1 in a form competent with peptidyl-tRNA cleavage ^49–51^. While structural studies ^43–45^ and single-molecule analysis ^52^ have revealed that eRF1 recognition of stop codons is highly specific and efficient, potential binding to near-cognate sense codons has been suggested ^53–55^. However, a genome-wide survey of eRF1-ribosome associations remains unaccomplished.

We reconciled these two distinct translation events; unexpectedly, our investigation of eRF1 codon recognition revealed endogenous ribosome pools rescued by the surveillance system in human cells. eRF1-selective Monosome-Seq and Disome-Seq showed that eRF1 binds not only to stop codons but also to near-cognate sequences, including UUA codons. This tentative UUA codon binding does not cause peptidyl-tRNA cleavage but leads to ribosome stalling and collision. Congruently, disomes predominantly accumulate on UUA codons upon double depletion of rescue factor ASCC3 and translation initiation repressor 4EHP, suggesting that the association of eRF1 with a UUA sense codon serves as a ribosome pause suppressed by the surveillance system in cells. Importantly, depletion of eRF1 by the degrader compound counteracted disome formation on UUA codons by ASCC3-4EHP deficiency, indicating the direct link between eRF1-mediated UUA stalling and the rescue. We also showed that an absence of ribosome collision repression by ASCC3-4EHP induces the stress response, including p38 phosphorylation, ATF3 expression, and transcriptome remodeling. Our study reveals natural ribosome rescue targets and the importance of this surveillance for maintaining cellular homeostasis.

## Results and discussion

### eRF1 binds to UUA sense codons and causes ribosome stalling

To determine the codon recognition diversity of eRF1, we isolated ribosomes associated with eRF1 and then subjected them to ribosome profiling (*i.e.*, selective Monosome-Seq) (Figure 1A) ^56,57^. For this purpose, we generated a stable HEK293 cell line that expressed streptavidin-binding peptide (SBP)-tagged eRF1 in a tetracycline-inducible manner (Figure S1A). Previous studies indicated that the N-terminal tagging, as we designed in this study, still maintains the functions of eRF1 ^52,55^. Simultaneous RNase treatment and immunoprecipitation (IP) allowed us to enrich ribosomes directly bound to eRF1 and exclude ribosomes and RNA-binding proteins (RBPs) bound through mRNAs in the form of polysomes (Figure 1A). Notably, to enable eRF1-baited pulldown of ribosomes and footprints, we used micrococcal nuclease (MNase), which maintains ribosome integrity ^58^, rather than RNase I, a standard RNase used for ribosome footprinting ^59–62^. Our MNase-based Monosome-Seq showed the typical size of ribosome footprints (Figure S1B) and the high reproducibility between replicates (Figure S1C). Compared to RNase I, MNase generates ribosome footprints with the fluctuating ends, due to its cleavage preference toward the A/U nucleotide ^63^. Nonetheless, our MNase-generated ribosome footprints still showed the 3-nt periodicity (Figure S1D-E), allowing the codon-resolution analysis. We determined the A-site offset for the footprints, considering the 5′ end of the footprints stemmed from ribosomes at stop codons (Figure S1F, typically 15 nt). These features were maintained in eRF1-selective Monosome-Seq (Figure S1B-F). Consistent with a report that eRF1 in the ribosomal A site protects footprints from 3′ end trimming by RNase ^64^, eRF1-selective Monosome-Seq showed a reduced amount of short (∼21-23 nt) footprints (Figure S1B).

**Figure 1.**
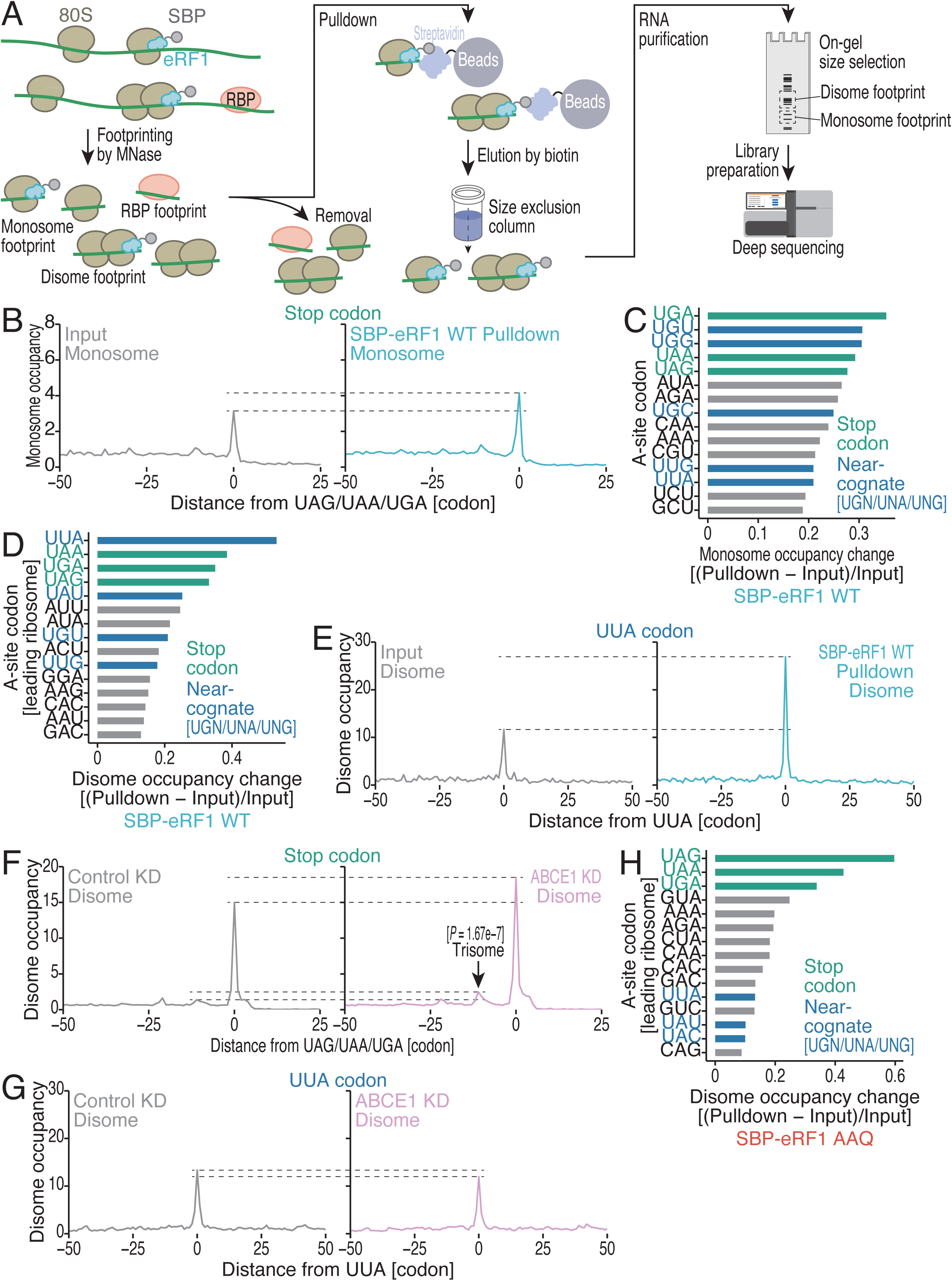
eRF1 transiently associates with UUA sense codons and leads to ribosome collision. (A) Schematic of eRF1-selective Monosome-Seq and Disome-Seq. (B) Metagene plot of the relative monosome footprint distribution (monosome occupancy) around stop codons. (C) Monosome occupancy differences between the eRF1 WT-bound fraction and the input. The data for the top 15 codons are shown. See Figure S1H for the full data. (D) Disome occupancy differences between the eRF1 WT-bound fraction and the input. The data for the top 15 codons are shown. See Figure S2E for the full data. (E) Metagene plot of the relative disome footprint distribution (disome occupancy; a site of the leading ribosome) around UUA codons. (F) Metagene plot of disome occupancy around stop codons for the indicated conditions. *P* value for the read enrichment at the trisome position was calculated with the Mann‒Whitney *U* test (two-tailed). (G) Metagene plot of disome occupancy around UUA codons for the indicated conditions. (H) Disome occupancy difference between the eRF1 AAQ-bound fraction and the input. The data for the top 15 codons are shown. See Figure S4E for the full data. See also Figures S1-4.

Compared with those from the input libraries, ribosome footprints from the eRF1-IP libraries were accumulated at stop codons (Figures 1B-C). However, eRF1 binding to mRNAs was not exclusive to stop codons; to our surprise, eRF1 was also associated with ribosomes at a subset of sense codons (Figures 1B-C and S1H) in a reproducible manner (Figure S1G). In particular, these sense codons included those that can be considered near-cognate stop codons, such as UGU, UGG, UGC, and UUA (Figures 1C and S1H), suggesting the possibility of misrecognition of the near-cognate codons by eRF1.

Given that eRF1-mediated sense codon misrecognition is unwelcome, we reasoned that this phenomenon may cause ribosome stalling and thus lead to a collision with trailing ribosomes. To test this possibility, we applied disome profiling ^36,38,41,65,66^ to the complexes isolated with SBP-eRF1 (*i.e.*, selective Disome-Seq) (Figures 1A and S2A), isolating 50–80 nt footprints (twice the length of monosome footprints). As observed in eRF1-selective Monosome-Seq experiments, a population of longer disome footprints was enriched in eRF1-selective Disome-Seq (Figure S2A). The heterogeneity of disome footprint lengths may stem from the conformational diversity of leading and trailing ribosomes, as discussed in a previous work ^36^. The disome footprints recovery was highly reproducible (Figure S2B). As in Mosome-Seq (Figure S1F), A-site offset (to leading ribosomes) in disome footprints was assigned via the leading ribosomes in disomes on stop codons (Figure S2C, 45 nt typically). Again, disomes containing eRF1 were enriched in stop codons (Figures 1D and S2D-E). Notably, we observed even stronger enrichment of disome footprints on near-cognate UUA codons than on cognate stop codons (Figure 1D and S2E).

Considering that UUA may be a part of a UAA or UAG stop codon via a −1 frameshift, we investigated the possibility that disomes formed on UUA codons could stem from stop codons translated along a −1 frame by frameshifting. However, the occupancy of eRF1-associated disomes on UUA-A and UUA-G was not greater than that on UUA-U (Figure S2F). Therefore, the stalling of eRF1-associated ribosomes on UUA is unlikely to be attributed to out-of-frame stop codons.

These data indicate that eRF1 recruitment to a UUA codon causes ribosome collision.

### UUA-misrecognizing eRF1 does not terminate the translation

Next, we investigated whether the association of eRF1 with a sense codon leads to a termination reaction or peptidyl-tRNA cleavage. After the termination reaction by eRF1, ABCE1 dissociates 60S subunits from 80S ribosomes on the stop codons for recycling. Since ABCE1 functions after peptidyl-tRNA cleavage, ribosome accumulation on stop codons by ABCE1 depletion ensures that proper translation termination reactions are induced at those positions ^67–71^. Here, we used Disome-Seq because of its high sensitivity for detecting ribosome stalling. Notably, the application of Ribo-FilterOut ^10^ techniques enabled greater recovery of usable reads than our previous approach involving Cas9-mediated removal of specific sequences (DASH: depletion of abundant sequences by hybridization) ^36,61^ (Figure S3A) because of the effective removal of contaminated rRNA fragments. Additionally, we utilized the Thor (T7 high-resolution original RNA) technique, which is based on T7 polymerase-mediated RNA amplification for low-input RNA ^72^. These modifications allowed stable library preparation for a limited number of disome footprints, as well as monosome footprints (Figure S3C-J).

Indeed, siRNA-mediated knockdown of ABCE1 (Figure S3B) increased the disomes with unrecycled ribosomes on stop codons (Figures 1F), as shown by the results of Monosome-Seq in earlier studies ^67–71^. Moreover, we observed an additional queue of collided ribosomes 10-codon (1 ribosome size) upstream from the stop codons (Figure 1F), indicating trisome formation ^36^. We found that disomes on UUA codons were not increased by ABCE1 depletion (Figure 1G), suggesting that ribosomes stalled on UUA codons do not result in termination reactions and subsequent recycling.

The same conclusion was supported by selective Monosome/Disome-Seq for peptidyl-tRNA cleavage-deficient eRF1. Conversion of the GGQ motif to AAQ allows a conformational change in eRF1, which is associated with GTP hydrolysis by eRF3, but it cannot induce the cleavage of peptidyl-tRNA ^43,45,49,73^. Ultimately, eRF1 AAQ may accumulate dead-end complexes of ribosomes after the accommodation of the middle (M) domain of eRF1 into the peptidyl transferase center (PTC) ^43,45^. While wild-type (WT) eRF1 proceeds to the accommodated state with subsequent ribosome recycling, AAQ eRF1 stacks in the accommodated state without further recycling (Figure S4A). Given that a significant fraction of ribosomes bound with eRF1 should be sequestered in the accommodated state in the AAQ mutant, relatively, the non-accommodated state in the pulldown should be decreased (Figure S4A). Employing this situation, we tested whether the eRF1-ribosome complex on UUA is in a non-accommodated state. As expected, the AAQ eRF1 mutant was associated with disomes in which the leading ribosomes are located on canonical stop codons (Figure 1H and Figure S4B-E). However, compared with WT eRF1, the association of AAQ eRF1 with disomes on UUA codons was attenuated (Figure 1D vs. 1H), indicating that eRF1 on the UUA codon does not proceed to the PTC accommodation step.

Thus, taking these data together, we conclude that eRF1 may bind tentatively to ribosomes on UUA codons but does not trigger the hydrolysis of peptidyl-tRNA.

### eRF1 delays ribosome traversal at UUA codons

Given these observations, we reasoned that misrecognition by eRF1 of UUA codons may compete with tRNA^Leu^_UAA_ and delay protein synthesis. To test this possibility, we harnessed an *in vitro* translation assay with rabbit reticulocyte lysate (RRL), which may have limited Leu-tRNAs due to high optimization for hemoglobin synthesis ^74,75^. We prepared two types of nanoluciferase (Nluc) reporter mRNAs: one predominantly uses CUG codons, the most frequent Leu codon in the human genome, and the other uses UUA codons. In total, 11 out of the 16 leucine codons in Nluc were substituted with CUG or UUA (Figure 2A, 11× CUG and 11× UUA). Upon the addition of recombinant eRF1-eRF3 to the reaction mixture, we observed decreased protein synthesis from the 11× UUA reporter but not from the 11× CUG reporter (Figure 2B).

**Figure 2.**
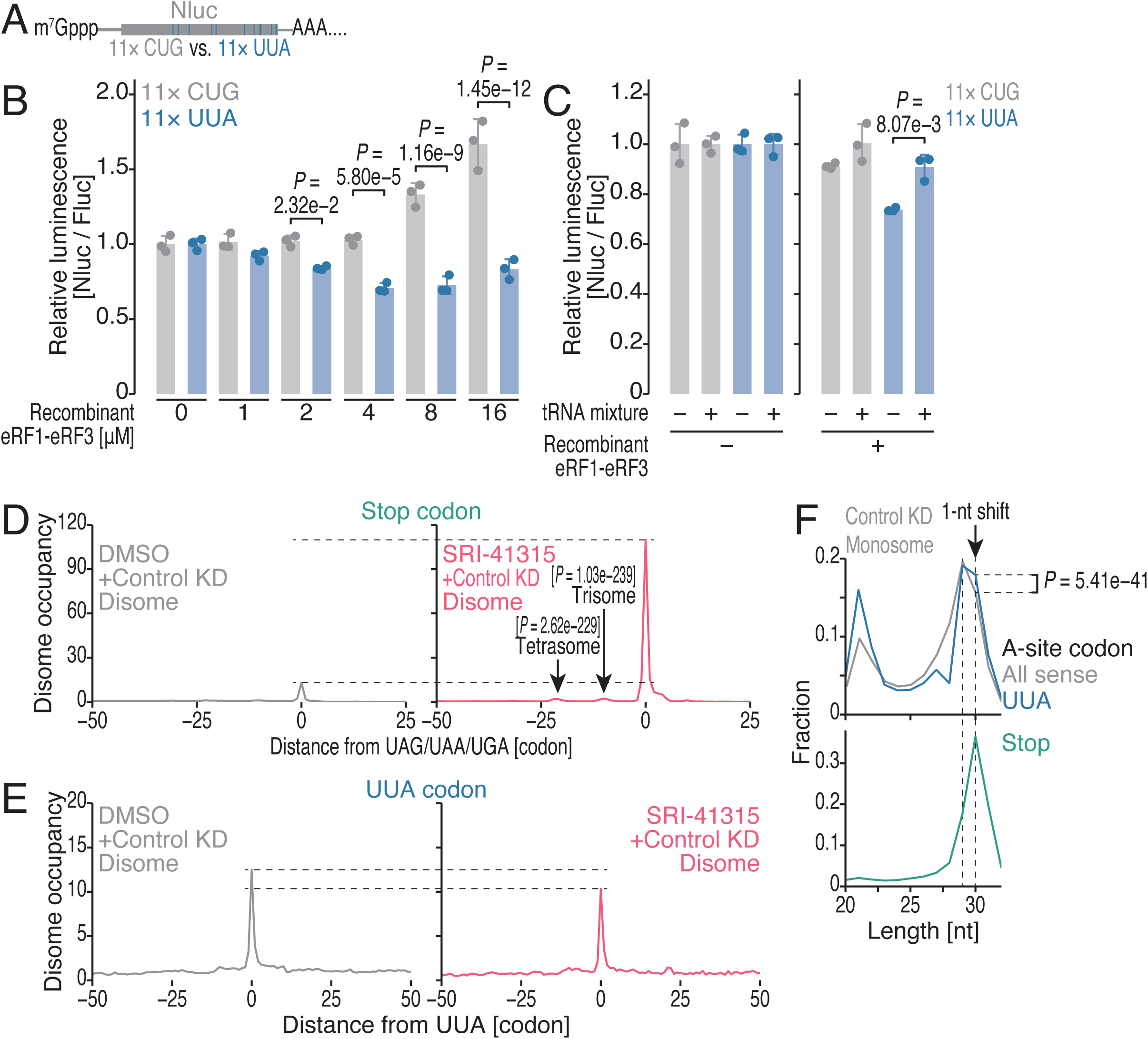
eRF1 delays ribosome elongation at UUA codons. (A) Schematic of reporter mRNAs encoding Nluc-11× CUG and 11× UUA. (B-C) *In vitro* translation assay using RRL for the reporter mRNAs shown in A. In B, a titrated amount of the recombinant eRF1-eRF3 complex was added to the reactions. In C, mammalian liver tRNA was further supplemented in the reaction with or without 2 μM recombinant eRF1-eRF3 complex. The data are presented as the means (bars) and s.d.s (errors) for replicates (points, n = 3). *P* values were calculated by the Tukey-Kramer test (two-tailed). (D) Metagene plot of disome occupancy around stop codons for the indicated conditions. *P* values for the read enrichment at the trisome position and the tetrasome position were calculated with the Mann‒Whitney *U* test (two-tailed). (E) Metagene plot of disome occupancy around UUA codons for the indicated conditions. (F) Distribution of monosome footprint length at the indicated codon species in the A site. *P* value for the enrichment of 30 nt footprint at UUA codons compared to that of all sense codons was calculated via Pearson’s chi-square test (two-sided). See also Figure S3.

Given that the supplementation of mammalian liver tRNA facilitates translation elongation on UUA codons in RRL ^75^, we reasoned that an additional tRNA pool counteracts eRF1-eRF3-mediated translation stalling on UUA codons. As expected, the translation repression by eRF1-eRF3 on the 11× UUA reporter was recovered when the tRNA mixture was added (Figure 2C).

To test that cellular eRF1 mediates ribosome stalling on UUA codons, we reduced eRF1 protein level by the degrader compound SRI-41315 ^55,76,77^ (Figure S5A) and then conducted Disome-Seq (Figure S5B-D). As expected, we observed huge disome buildup on stop codons and associated trisome and tetrasome upstream (Figure 2D). In stark contrast, disomes on UUA codons were rather reduced by the loss of eRF1 (Figure 2E). We note that the disome reduction by SRI-41315 is codon specific but not global (Figure S5E). These data support that eRF1 frequently misrecognizes UUA codons in cells, leading to substantial ribosome stalling.

Moreover, we observed that eRF1 binding to UUA codons is a stoichiometric event. Stop codon recognition by eRF1 pulls the 4th position nucleotide into the ribosome A site ^43,44^, allowing the 1-nt longer ribosome footprints ^78^. Consistent with these reports, we observed 30-nt footprints, 1-nt longer than major 29-nt footprints, at stop codons (Figure 2F, bottom). We found that footprints at the UUA codons showed an increased population of 30-nt footprints (Figure 2F, top), suggesting that a substantial fraction of ribosomes on UUA were associated with eRF1. In contrast, we did not observe such a footprint extension on the CUG codon (Figure S3K).

These data indicate that the misrecognition of UUA by eRF1 leads to a delay in elongation and subsequent ribosome collision.

### mRNA surveillance system suppresses ribosome collisions formed on UUA codons

The ribosome collision induced by sense codon-misrecognizing eRF1 led us to investigate the involvement of the ribosome rescue processes. For this purpose, we surveyed disomes accumulated due to the loss of key factors that resolve/suppress disomes. Recent findings have demonstrated that the leading ribosome in a ribosome collision can be split by the ATP-dependent helicase ASCC3 ^13,17,18,21–23^ (Figure 3A). Moreover, colliding ribosomes can also act as platforms to repress translation initiation, preventing further ribosome collision, through the recruitment of 4EHP, a cap-binding protein ^24–28^ (Figure 3A). We knocked down these genes to maximize the generation of disomes that are subjected to the surveillance system (Figures S3B and S5A).

**Figure 3.**
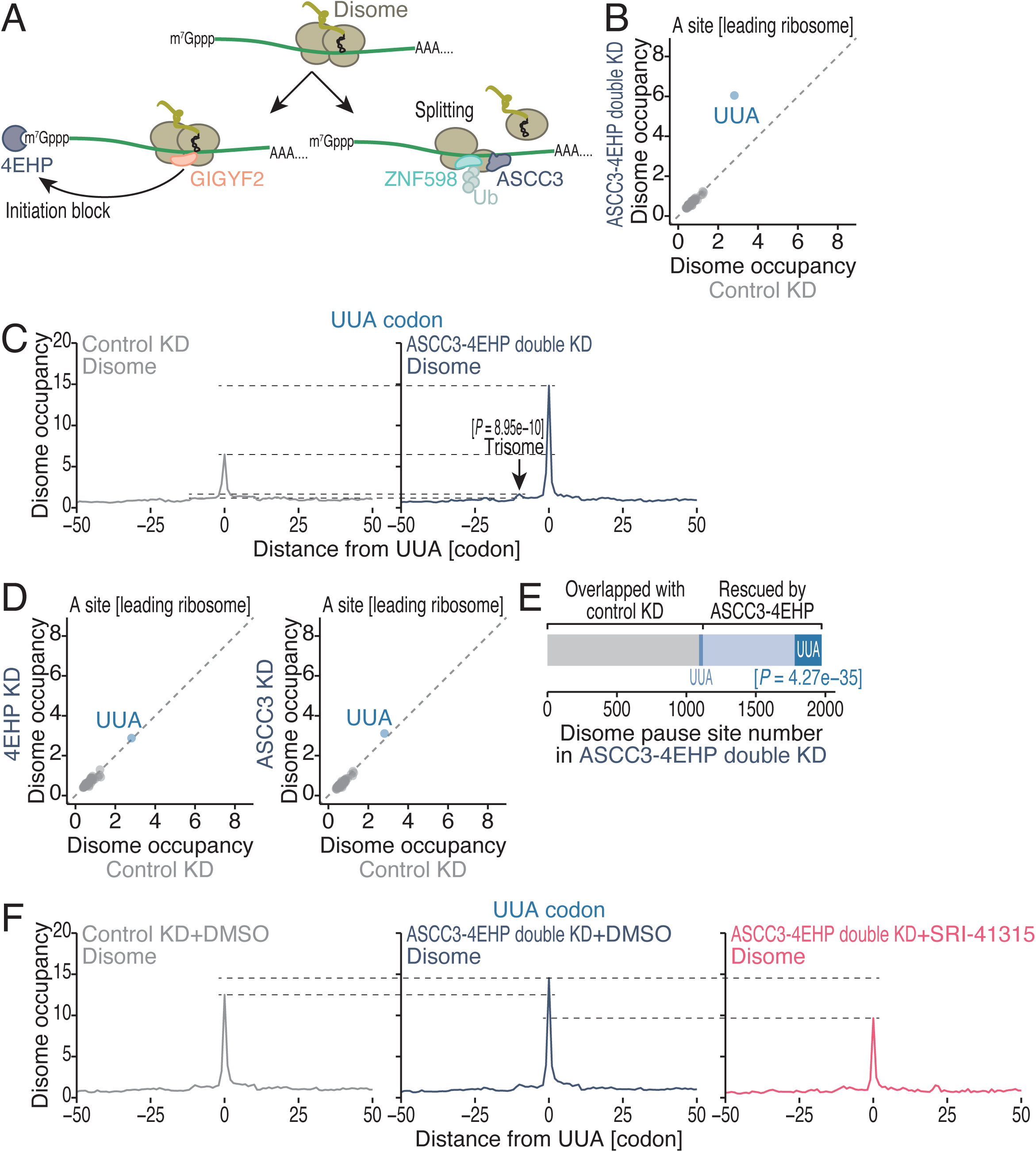
Disomes on UUA codons are rescued by the surveillance system. (A) Schematic of the translation initiation block and ribosome splitting associated with ribosome collision. (B and D) Comparison of disome occupancy on A-site codons (for the leading ribosomes) for the indicated conditions. Data for sense codons are shown. (C and F) Metagene plot of disome occupancy around UUA codons for the indicated conditions. *P* value for read enrichment at the trisome position was calculated with the Mann‒Whitney *U* test (two-tailed). (E) A breakdown of disome pause sites, illustrating the proportions of the indicated categories. *P* values were calculated via Pearson’s chi-square test (two-sided). See also Figures S3 and S5 and Tables S1-5.

Remarkably, upon ASCC3-4EHP double depletion, we observed a pile-up of disomes where leading ribosomes were at UUA codons (Figures 3B and S3C-J, and Tables S1 and S3). These collisions at UUA codons subsequently led to queued trisomes (Figure 3C). On the other hand, a single knockdown of each factor showed limited effect on ribosome collision at UUA codons (Figure 3D and Tables S4 and S5). This suggests that multiple regulatory points may synergistically function to resolve/suppress ribosome collision.

We note that, even without ASCC3-4EHP depletion, UUA codons were disome-prone context (Figure 3B-D), consistent with earlier work ^36^. The UUA codon is rare in the human genome, and the corresponding tRNA^Leu^_UAA_ was expressed at a low level in HEK293T cells ^79^ (Figure S6A). However, the rare codons and low-level tRNAs did not always lead to the ribosome collision (Figure S6A), highlighting the uniqueness of the UUA codon.

In addition to the general trends involving UUA codons, our data revealed individual ribosome collision sites that are rescued by the system. In the ASCC3-4EHP double-depleted cells, we identified 1973 disome peak sites (see Methods), ∼60% of which were already present in the control knockdown samples (Figure 3E). The remaining ∼40% (854 sites) represented ribosome collision sites that emerged upon the loss of ASCC3-4EHP. Again, the newly generated disome pause sites were found at UUA codons at a statistically significant level (Figure 3E), which is consistent with the global increase in high disome occupancy on UUA codons (Figure 3B). These loci were exemplified by UUA codons in *PNRC1* and *CCT7* mRNAs (Figure S6B-C).

Notably, unlike those in yeast ^38,39^, the poly-Lys motifs found in endogenous mRNAs were not subjected to ribosome rescue under our conditions. Although the Lys-rich motif of *MTDH* mRNA is known to induce ribosome stalling ^36,80^, ribosome collision at this motif was not further strengthened by ASCC3-4EHP double depletion (Figure S6D). More broadly, the defect in surveillance system had limited effects on disome occupancy at Lys-Lys motifs (P-A sites) (Figure S6E). This observation was congruent with an earlier report that the human ribosome rescue system recognizes longer Lys-AAA repeats than the yeast system does ^13^. The human genome tends to evade such long repeats.

Given the correspondence of disome formation at UUA codons, we hypothesized that ribosome collision induced by eRF1 misrecognition of UUA may generate substrates for ribosome rescue in cells. To test this scenario, we further depleted eRF1 by SRI-41315 treatment from ASCC3-4EHP double knockdown cells (Figure S5A) and performed Disome-Seq (Figure S5B-D). Strikingly, the reduction of eRF1 suppressed ribosome collision evoked by ASCC3-4EHP loss (Figure 3F). We note that relatively higher UUA disome in control knockdown (Figure 3F vs. 3C) may be due to the DMSO solvent ^81^. Again, UUA codons in *PNRC1* and *CCT7* served as representative examples (Figure S6F-G).

Our list of ribosome collision sites in the factor knockdown also showed a functional difference between ABCE1 and ASCC3-4EHP. Although ABCE1 knockdown generated new disome pause sites (Figure S6H and Table S2), the overlap with ones found in ASCC3-4EHP double depletion was limited (Figure S6I).

These data together demonstrate that eRF1 recruitment to UUA codons is a cause of ribosome stalling that can be targeted by the surveillance mechanism.

### Sustained ribosome collisions at UUA codons trigger a stress response and ATF3 expression

Next, we investigated the cellular responses triggered when colliding ribosomes were left unresolved at UUA codons. In addition to the RQC pathway, other responses, including the ribotoxic stress response (RSR) mediated by MAP3K ZAKα activation, have been highlighted in recent studies ^82–89^. In the RSR, ZAKα activates downstream signaling pathways, including p38 activation ^82–89^. We reasoned that the accumulation of disomes upon depletion of ASCC3 and 4EHP may trigger crosstalk with the RSR. Consistent with this scenario, we observed the phosphorylation of p38, a hallmark of p38 activation (Figure 4A).

**Figure 4.**
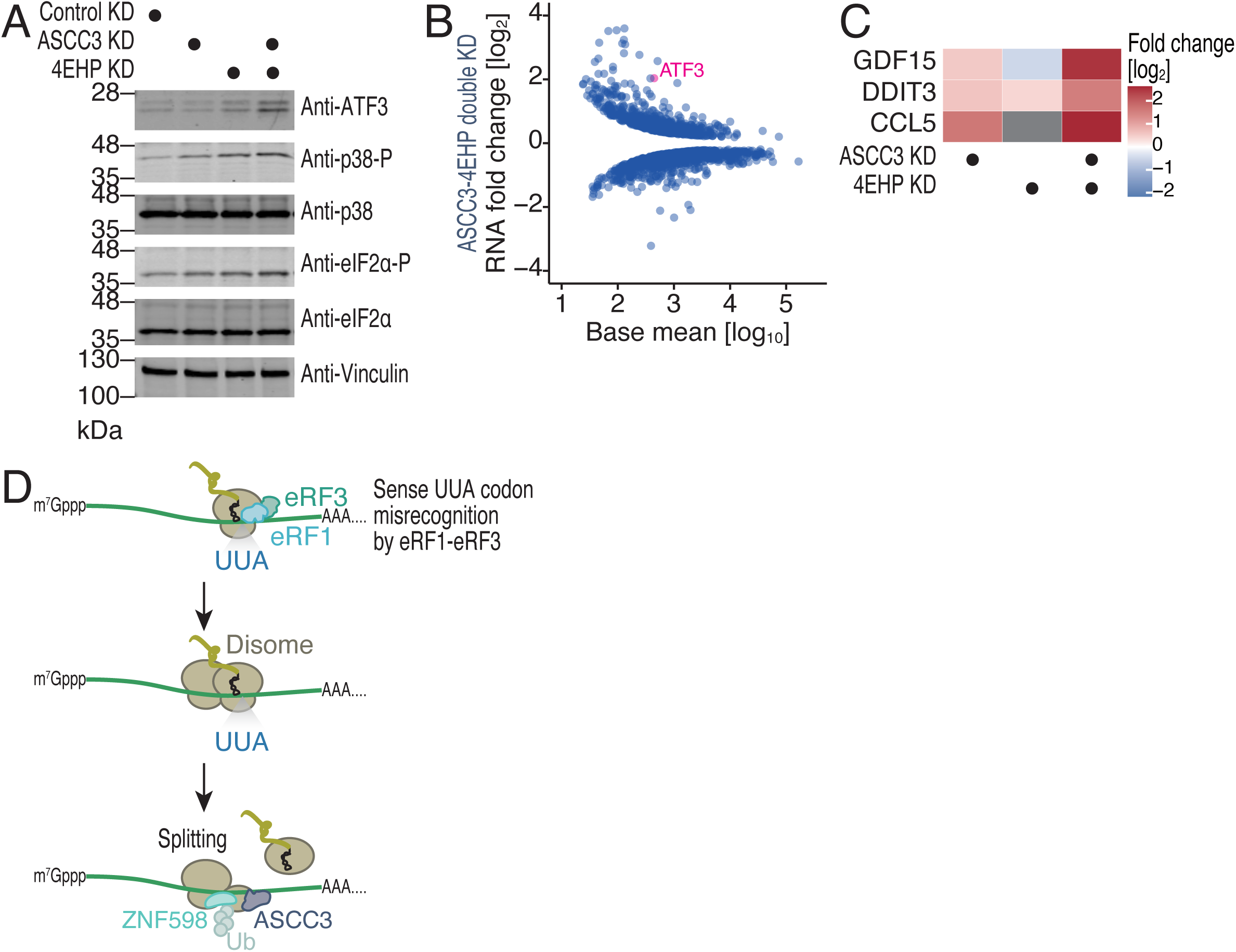
Depletion of ASCC3 and 4EHP activates alternative pathways that sense disomes. (A) Western blotting of the indicated proteins. Vinculin was used as a loading control. Representative data from three replicate experiments are shown. (B) MA (M, log ratio; A, mean average) plot for RNA abundance change, measured by RNA-Seq, with ASCC3-4EHP double knockdown. Transcripts with significant changes (FDR < 0.05) are shown. (C) Heatmap of RNA abundance change for the indicated genes under the indicated conditions. The color scales for the fold changes are shown. (D) Schematic of the model of eRF1-mediated UUA codon association, ribosome collision, and subsequent rescue by ASCC3-4EHP. See also Figure S6.

A recent study showed that RSR activation induces transcriptome remodeling through the accumulation of the stress-inducible transcription factor ATF3 ^90^. Through RNA-Seq (Figure S6J), we found that ATF3 expression was increased upon the double knockdown of ASCC3 and 4EHP (Figure 4A). The accumulation of the ATF3 protein was attributed to the upregulation of its mRNA abundance (Figure 4B), along with large-scale alterations in the cellular transcriptome. The double depletion of ASCC3 and 4EHP induced the expression of direct targets of ATF3 (such as GDF15, DDIT3, and CCL5) ^91^ (Figure 4C). These data suggest that unresolved disomes activate stress response to induce cellular transcriptome remodeling.

## Conclusion

Our work reveals unexpected impacts of the association of eRF1 with the UUA sense codon for translation quality control (Figure 4D). Recent work ^92^ has shown that Leu depletion leads to pronounced ribosome stalling specifically at the UUA codon among the six Leu codons. This heightened sensitivity may arise, at least in part, from competition between the UUA-decoding tRNA^Leu^_UAA_ and eRF1 at the A site. The ability of eRF1 to engage rare, non-optimal codons has been reported in *Neurospora*, *Drosophila*, and yeasts ^53,54^. Moreover, a molecular glue (such as SRI-41315) between eRF1 and ribosomes may induce recognition of near-cognate UYR codons (where Y represents U or C and R represents A or G), which includes the UUA ^55^. Although in these artificial setups, eRF1 ultimately terminates translation on those cryptic sites ^53–55^, cellular eRF1 does not proceed to peptidyl-tRNA hydrolysis. Our results show that eRF1 may transiently associate with widespread sense codons before full accommodation by GTP hydrolysis of eRF3 in cells. Near-cognate UUA codons may extend the dwell time of eRF1-eRF3, slow ribosome traversal by competing with cognate tRNAs, induce the unwelcome ribosome collision, and evoke ribosome rescue. eRF1-mediated translation termination and ASCC3-mediated rescue both generated truncated proteins. However, the latter mechanism is safer, since it is associated with nascent protein degradation to avoid the accumulation of harmful protein. The human genome preserves the rarity of UAA codon usage, which circumvents the natural mRNA surveillance events.

## Limitations of this study

After ribosome rescue, peptidyl-tRNA associated with the 60S subunit is subjected to CAT-tailing and subsequent degradation ^1–4^. In the present study, we did not examine the fate of nascent chains synthesized by ribosomes rescued via ASCC3. In addition, stall-prone mRNAs may be targeted by no-go decay (NGD), which degrades problematic transcripts ^20,93,94^ and shares ZNF598 as its trigger. Recently, endogenous mRNAs subject to NGD have been systematically surveyed in vertebrates ^95^. Future studies will be needed to elucidate the potential coordination between ribosome rescue pathways and NGD.

## Acknowledgments

We are grateful to all the members of the Iwasaki laboratory and Dr. Mariko Sasaki for their constructive discussions and technical help. Computation was supported by the HOKUSAI SailingShip supercomputer facility at RIKEN. We are grateful to the Support Unit for Bio-Material Analysis, RIKEN CBS Research Resources Division, for Sanger sequencing. We also thank Dr. Nicholas T. Ingolia for sharing plasmids. This work was supported by the Ministry of Education, Culture, Sports, Science and Technology (MEXT) (JP20H05784 and JP24H02307 to S.I.; JP21H05281 to T.I.), the Japan Agency for Medical Research and Development (AMED) (JP20gm1410001 to S.I. and T.I.), the Japan Society for the Promotion of Science (JSPS) (JP23H02415 and JP23H00095 to S.I.; JP22J13110 and JP22KJ0894 to P.H.), and RIKEN (Pioneering Project to S.I. and T.I.; Incentive Research Project to A.Y.). P.H. was a JSPS Research Fellow (DC2/PD).

## Author contributions

Conceptualization: P.H. and S.I.; Methodology: P.H., M.M., A.Y., and S.I.;

Formal analysis: P.H., M.M., A.Y., and T.I.;

Investigation: P.H., M.M., A.Y., and T.I.;

Resources: T.I.;

Writing – Original Draft: P.H. and S.I.;

Writing – Review & Editing: P.H., M.M., A.Y., T.I., and S.I.; Visualization: P.H., A.Y., and S.I.;

Supervision: T.I. and S.I.;

Project administration: S.I.;

Funding Acquisition: P.H., A.Y., T.I., and S.I.

## Methods

### Cell lines and gene knockdown

HEK293 Flp-In T-REx cells (Thermo Fisher Scientific) were maintained at 5% CO_2_ and 37°C in DMEM with high glucose and GlutaMAX supplement (Thermo Fisher Scientific) containing 10% fetal bovine serum (FBS). The cell cultures were routinely tested for contamination with *Mycoplasma* spp. via an e-Myco VALiD Mycoplasma PCR Detection Kit (iNtRON Biotechnology) and were confirmed to be free of infection.

Stable cell lines expressing SBP-eRF1 WT or SBP-eRF1 AAQ mutant were generated (see the “*DNA construction*” section below) via cotransfection of the expression plasmids with pOG44 (Thermo Fisher Scientific) by X-tremeGENE 9 (Sigma–Aldrich) and selected via the addition of hygromycin B (InvivoGen) according to the manufacturer’s recommendation.

Gene knockdown was conducted in HEK293 Flp-In T-REx cells via the transfection of siRNAs (ON-TARGETplus/siGENOME SMARTpool, Dharmacon) by TransIT-X2 Dynamic Delivery System (Mirus) according to the manufacturer’s recommendation. This study used the following siRNAs: control, D-001810-10-50; ABCE1, L-008702-00-0020; ASCC3, L-012757-01-0010; 4EHP, L-019870-01-0010; and eRF1, M-019840-01-0005. For Thor-Monosome-Seq, Thor-Disome-Seq, and Western blotting, siRNA transfection was conducted twice before the cells were harvested. During the first transfection, a total concentration of 75 nM siRNA was transfected; 50 nM ABCE1 siRNA, 25 nM ASCC3 siRNA, and 25 nM 4EHP siRNA were adjusted with control siRNA to render 75 nM siRNA in total. After 72 h of incubation and cell reseeding, the second siRNA was used at a dose of 30 nM. The cells were incubated for another 24 h and harvested.

For the experiments involving eRF1 depletion, gene knockdown was conducted for 48 h as described above. At 4 h before harvest, cells were treated with 10 μM SRI-41315 (Sigma–Aldrich).

### DNA construction

#### pCDNA5/FRT/TO-SBP-eRF1 WT and AAQ

The DNA fragment encoding eRF1 was PCR-amplified from cDNA generated with total RNA from HEK293 Flp-In T-REx cells as a template and inserted into pCDNA5/FRT/TO-SBP-eIF4A1 ^96^, replacing eIFA1 with eRF1. AAQ substitution was introduced via site-directed mutagenesis.

#### psiCHECK2 Nluc-11× CUG, Nluc-11× UUA, and Fluc

Nluc-11× CUG was based on Nluc encoded in pNL1.1 (Promega); Leu24-CUU and Leu32-UUG were converted to CUGs. The synthesized DNA (Eurofins) was inserted into psiCHECK2-GPX1 ^96^, replacing the *Renilla* luciferase (RLuc)-HSV TK promoter-Fluc region with Nluc-11× CUG and placing the V5 tag at the N-terminus of Nluc-11× CUG.

In Nluc-11× UUA, the CUG codons in Nluc-11× CUG were replaced with UUA codons. The synthesized DNA (Eurofins) was subsequently cloned as described above.

Similarly, DNA fragments encoding Fluc were PCR-amplified from psiCHECK2-GPX1 as a template and subsequently cloned as described above.

### eRF1-selective Monosome-Seq and Disome-Seq

Protein expression from stable integrants of SBP-eRF1 WT or AAQ in HEK293 Flp-In T-REx cells was induced with 0.2 μg/ml tetracycline for 2 d before cell harvest. Then, the cells were washed with ice-cold PBS once and lysed with 400 μl of lysis buffer (20 mM Tris-HCl pH 7.5, 50 mM NaCl, 5 mM MgCl_2_, 1 mM DTT, 1% Triton X-100, and 100 μg/ml cycloheximide). The cell lysates were treated with 0.0375 U/μl TURBO DNase (Thermo Fisher Scientific) on ice for 10 min and then clarified by centrifugation at 20,000 × *g* at 4°C for 10 min. The supernatants were aliquoted, flash-frozen in liquid nitrogen, and stored at −80°C. RNA concentrations of the cell lysates were determined by a Qubit RNA BR Assay kit (Thermo Fisher Scientific).

The cell lysate containing 6 μg of total RNA was incubated with 70 μl of Dynabeads M-280 Streptavidin (Thermo Fisher Scientific), which was equilibrated with lysis buffer, at 4°C for 40 min. The mixture was treated with 3000 U MNase (TaKaRa) with 5 mM CaCl_2_ at 25°C for 45 min. A fraction of the reaction mixture was used as the input sample.

The reaction mixture was placed on a magnetic stand, and the beads were washed six times with wash buffer (20 mM Tris-HCl pH 7.5, 150 mM NaCl, 5 mM MgCl_2_, 1 mM DTT, 1% Triton X-100, and 100 μg/ml cycloheximide). Ribosomes bound to SBP-eRF1 were eluted with a wash buffer containing 5 mM biotin. For the input sample after RNase digestion, ribosome complexes were isolated with MicroSpin S-400 HR columns (Cytiva)^61^. RNA was isolated with TRIzol LS (Thermo Fisher Scientific) and a Direct-zol RNA Microprep Kit (Zymo Research).

After PAGE using 15% denatured polyacrylamide SuperSep RNA gels (Fujifilm Wako Pure Chemical Corporation), RNA fragments within the range of 17–34 nt (for Monosome-Seq) and 50–80 nt (for Disome-Seq) were gel-excised ^61^. After dephosphorylation, linker ligation, rRNA depletion via a stand-alone Ribo-Zero Gold rRNA Removal Kit (Human/Mouse/Rat) (Illumina), reverse transcription, circularization, and PCR amplification ^61^, the DNA libraries were sequenced via a NovaSeq 6000 (Illumina) with the paired-end/150 nt-long read option.

### Thor-Monosome-Seq and Thor-Disome-Seq

siRNA-mediated gene knockdown was performed as described in the “*Cell lines and gene knockdown*” section above. The cells were washed with ice-cold PBS once and then lysed with 400 μl of modified lysis buffer (20 mM Tris-HCl pH 7.5, 150 mM NaCl, 5 mM MgCl_2_, 1 mM DTT, 1% Triton X-100, 100 μg/ml cycloheximide, 100 μg/ml tigecycline, and 100 μg/ml chloramphenicol). The cell lysates were treated with 0.025 U/μl TURBO DNase (Thermo Fisher Scientific) on ice for 10 min and then clarified by centrifugation at 20,000 × *g* at 4°C for 10 min. The supernatants were aliquoted, flash-frozen in liquid nitrogen, and stored at −80°C. RNA concentration of the cell lysates was determined by a Qubit RNA BR Assay kit (Thermo Fisher Scientific).

The cell lysate containing 3 μg of total RNA was adjusted to 300 μl with modified lysis buffer and treated with 0.067 U/μl RNase I (LGC Biosearch Technologies) at 25°C for 45 min. The reaction was stopped by transferring the mixture to ice, followed by the addition of 200 U SUPERase•In RNase Inhibitor (Thermo Fisher Scientific). The mixture was layered over 900 μl of 1 M sucrose cushion ^61^ and ultracentrifuged at 100,000 rpm (543,000 *× g*) at 4°C for 1 h using a TLA-110 rotor and an Optima MAX-TL Ultracentrifuge (both Beckman Coulter).

Then, Ribo-FilterOut was conducted ^10^. After the supernatant was carefully removed, the ribosome pellet was recovered via the addition of 150 μl pellet lysis buffer (20 mM Tris-HCl pH 7.5, 300 mM NaCl, 5 mM EDTA, 1 mM DTT, 1% Triton X-100, and 0.02 U/μl SUPERase•In RNase Inhibitor) and incubated on ice for 10 min. Ribosome footprints were further purified by centrifugation at 14,000 × *g* at 4°C for 10 min in a 0.5 ml Amicon Ultra 100 kDa cutoff centrifugal filter unit (Merck Millipore). From the flow-through, RNA was isolated with TRIzol LS (Thermo Fisher Scientific) and a Direct-zol RNA Microprep Kit (Zymo Research). After PAGE using 15% denatured polyacrylamide SuperSep RNA gels (Fujifilm Wako Pure Chemical Corporation), RNA fragments within the range of 17–34 nt (for Monosome-Seq) and 50–80 nt (for Disome-Seq) were gel-extracted ^61^.

We employed the Thor approach ^72^ for both Monosome-Seq and Disome-Seq. After dephosphorylation, linker ligation, and rRNA depletion by Ribo-Zero, accompanied by a TruSeq Stranded Total RNA Kit (Illumina) or riboPOOL human Ribo-seq (siTOOLs Biotech), RNA fragments were amplified via RNA-templated transcription using a T7-Scribe Standard RNA IVT Kit (CELLSCRIPT) for 2.5 h at 37°C ^72^. After the second linker was ligated to the 3′ end of the RNA by T4 RNA Ligase 2, truncated KQ [New England Biolabs (NEB)], cDNA was synthesized using ProtoScript II Reverse Transcriptase (NEB). DNA libraries were generated via PCR with Phusion High-Fidelity DNA Polymerase (NEB), adding barcodes for the Illumina sequencing platform, and sequenced via a NovaSeq 6000 or NovaSeq X Plus (Illumina) with the paired-end/150 nt-long read option.

### RNA-Seq

Total RNA was isolated from the cell lysates used for Thor-Monosome-Seq and Thor-Disome-Seq with TRIzol LS (Thermo Fisher Scientific) and a Direct-zol RNA Microprep kit (Zymo Research). Libraries were constructed using a TruSeq Stranded Total RNA Library Prep Kit (Illumina) and sequenced with a NovaSeq 6000 (Illumina) with the paired-end/150 nt-long read option.

### Deep sequencing data analysis

#### RNA-Seq

Using fastp (ver. 0.21.0) ^97^, the adaptor sequence was trimmed from the raw fastq. Read 1 from the pair-end sequencing library was used for downstream analysis. Reads that were aligned to noncoding RNA by STAR (ver. 2.7.0a) ^98^ were excluded. The remaining reads were aligned to the hg38 genome (GENCODE v32 annotation) via STAR (ver. 2.7.0a) ^98^. Duplicated reads that occurred during PCR were suppressed by UMI via UMI-tools (ver. 1.1.4) ^99^. Mapped reads were counted, and differential RNA expression was analysed using DESeq2 (ver. 1.28.1) ^100^.

#### Ribo-Seq and Disome-Seq

Using fastp (ver. 0.21.0) ^97^, low-quality reads in read 1 were corrected by read 2 of the pair-end sequencing data, and the adaptor sequence was trimmed from the raw fastq of read 1. Reads that were aligned to noncoding RNA by STAR (ver. 2.7.0a) ^98^ were excluded. The remaining reads were aligned to the hg38 genome (GENCODE ver. 32 annotation) via STAR (ver. 2.7.0a) ^98^. Duplicated reads that occurred during PCR or *in vitro* transcription by T7 polymerase were suppressed by UMI via UMI-tools (ver. 1.1.4) ^99^.

Footprints on stop codons from all genes in monosome or disome libraries were used to determine the distance from the 5′ end of the footprints to the A site of the ribosome or the leading ribosome in a disome.

Ribosome footprints mapped to the canonical mRNA transcripts (GENCODE ver. 32) were used to calculate average ribosome occupancy at any codon across the genome. Monosome and disome occupancies at any position (codon) were defined as the ratio between raw footprint read count and the average footprint read count per codon of the ORF. Transcripts that had fewer than 0.2 footprints per codon were excluded from the analysis. Codons that had fewer than two mapped reads were also excluded. Codons whose disome occupancy was the value of the mean + s.d. or larger were defined as disome pause sites.

Published tRNA-Seq data (GEO: GSE66550) ^79^ were processed as described previously ^36^. Codon usage frequency was calculated as described previously ^101^.

### Western blotting

The concentration of total protein in the cell lysates was adjusted. After 2× SDS sample buffer (125 mM Tris-HCl pH 6.8, 20% glycerol, 4% SDS, 10% 2-mercaptoethanol, and 0.004% BPB) was added, the samples were heated at 95°C for 5 min. Electrophoresis of the protein samples was performed via 5‒20% Tris‒glycine SDS‒PAGE gels (Gellex). Proteins were subsequently transferred to 0.2-μm nitrocellulose membranes via a Trans-Blot Turbo Transfer System (Bio-Rad). The membranes were blocked with TBS blocking buffer (LI-COR Biosciences) for 1 h at room temperature.

The primary antibodies used were anti-ABCE1 (Abcam, ab185548, 1:1000), anti-ASCC3 (PROTEINTECH, 17627-1-AP, 1:500), anti-4EHP (Cell Signaling Technology, 6916S, 1:250), anti-4EHP (Cell Signaling Technology, 79940S, 1:500), anti-eRF1 (Cell Signaling Technology, 13916S, 1:1000), anti-Vinculin (LI-COR Biosciences, 926-42215, 1:1000), anti-SBP Tag (Santa Cruz, sc-101595, 1:1000), anti-phospho-eIF2α (Cell Signaling Technology, 3398S, 1:1000), anti-eIF2α (Cell Signaling Technology, 9722S, 1:1000), anti-p38 MAPK (Cell Signaling Technology, 9212S, 1:1000), anti-phospho-p38 MAPK (Cell Signaling Technology, 4511S, 1:1000), anti-ATF3 (Cell Signaling Technology, 33593S, 1:1000), and anti-β-actin (LI-COR Biosciences, 926-42210, 1:1000). The following secondary antibodies were used: IRDye 800CW anti-rabbit IgG (LI-COR Biosciences, #926-32211, 1:10000) and IRDye 800CW anti-mouse IgG (LI-COR Biosciences, #926-32210, 1:10000). The membranes were imaged via an ODYSSEY CLx system (LI-COR Biosciences).

### Preparation of reporter mRNAs

The DNA fragments encoding the Nuc and Fluc reporters were PCR-amplified with the primers 5′-TGACTAATACGACTCACTATAGG-3′ and 5′-TGTATCTTATCATGTCTGCTCGAAG-3′ using psiCHECK2-Nluc-11× CUG, 11× UUA, and Fluc as templates. Using these DNA fragments as templates, *in vitro* transcription was conducted with a T7-Scribe Standard RNA IVT Kit (CELLSCRIPT) for 2 h at 37°C. Capping and polyadenylation were conducted using a ScriptCap m^7^G Capping Kit (CELLSCRIPT), a ScriptCap 2′-*O*-Methyltransferase Kit (CELLSCRIPT), and an A-Plus Poly (A) Polymerase Tailing Kit (CELLSCRIPT).

### Reporter assay with an in vitro translation system

*In vitro* translation of the reporter mRNAs (0.125 μM Fluc mRNA and 0.25 μM Nluc mRNA) was performed with Rabbit Reticulocyte Lysate [Nuclease-Treated] (Promega) at 30°C for 30 min. The indicated amount of eRF1-eRF3 recombinant proteins, which were prepared as previously described ^102^, was added to the reactions. For the indicated groups, mammalian liver tRNA (Sigma–Aldrich) was included at a final concentration of 0.2 mg/ml. The reaction was stopped by adding 1× Passive Lysis Buffer (Promega). A dual-luciferase reporter assay was performed via a Nano-Glo Dual-Luciferase Reporter Assay System (Promega) and a GloMax Navigator Microplate Luminometer (Promega).

## Data availability

The Monosome-Seq, Disome-Seq, and RNA-Seq data (GSE275336) [https://www.ncbi.nlm.nih.gov/geo/query/acc.cgi?acc=GSE275336] obtained in this study were deposited in the National Center for Biotechnology Information (NCBI) database. Gene annotations (GENCODE Human Release 32 reference) were obtained via the UCSC Genome Browser (https://genome.ucsc.edu/index.html).

## Code availability

The custom scripts used in this study are available at Zenodo (DOI: 10.5281/zenodo.13352233) [https://zenodo.org/doi/10.5281/zenodo.13352233].

---

**Figure S1.**
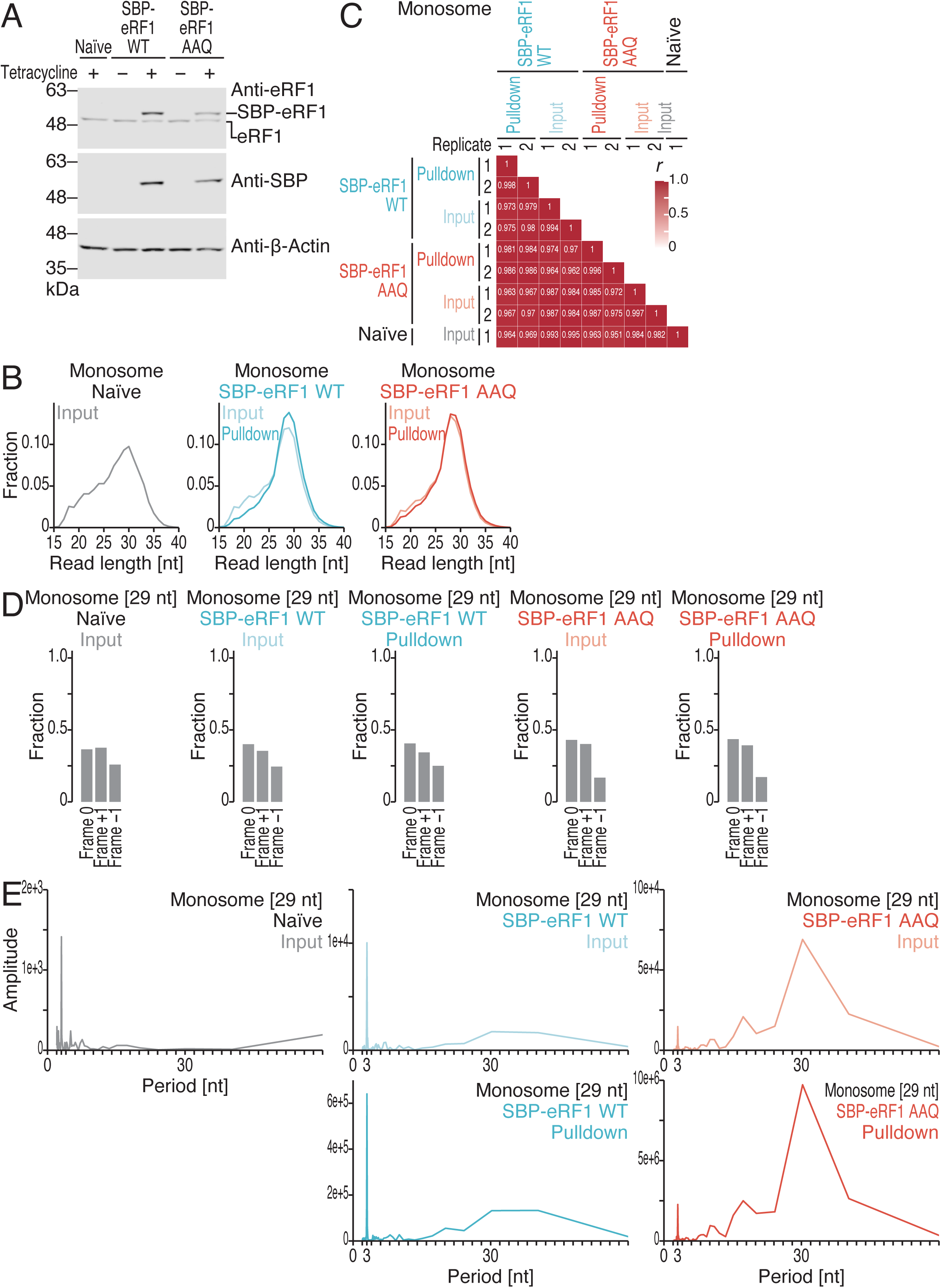
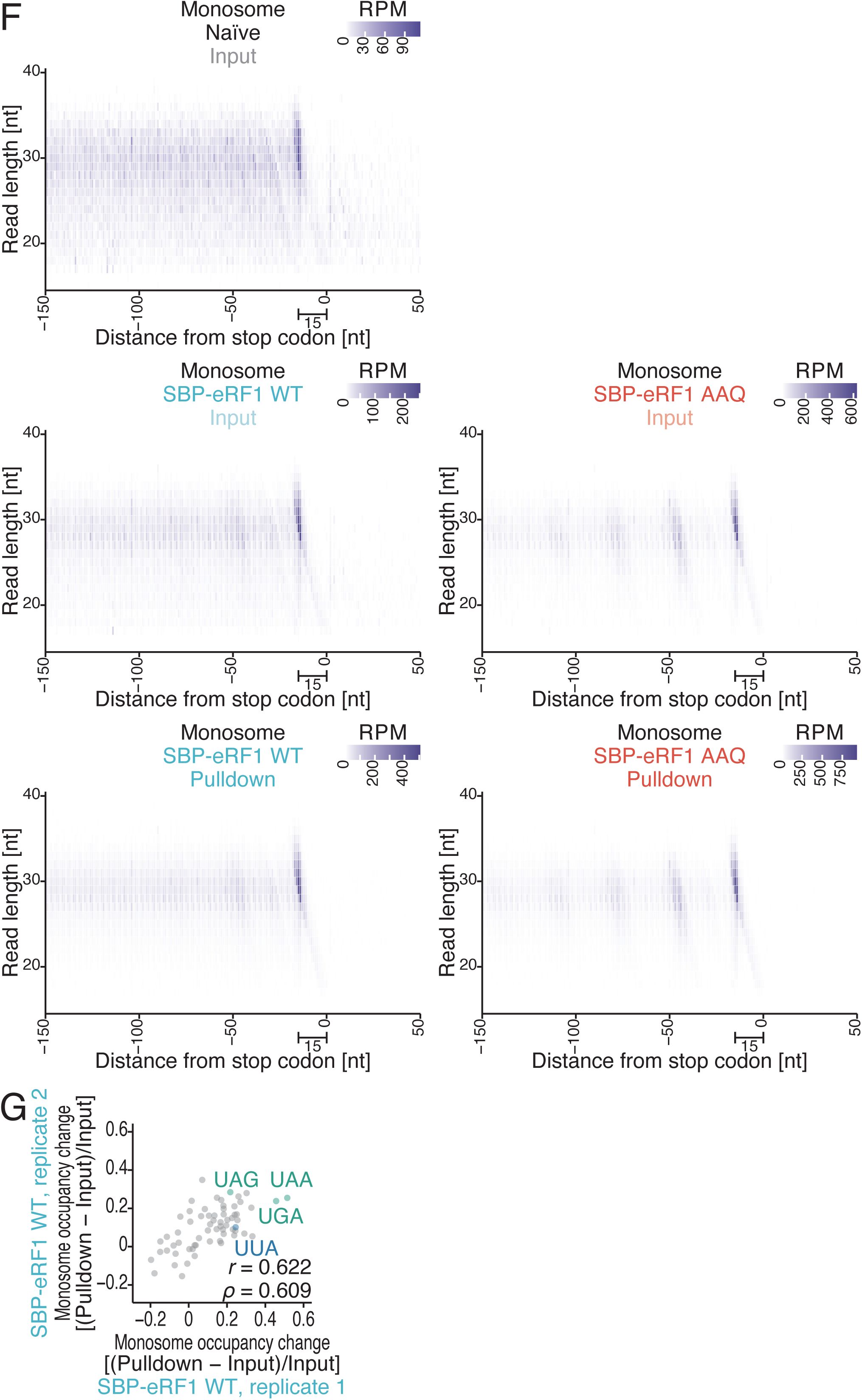
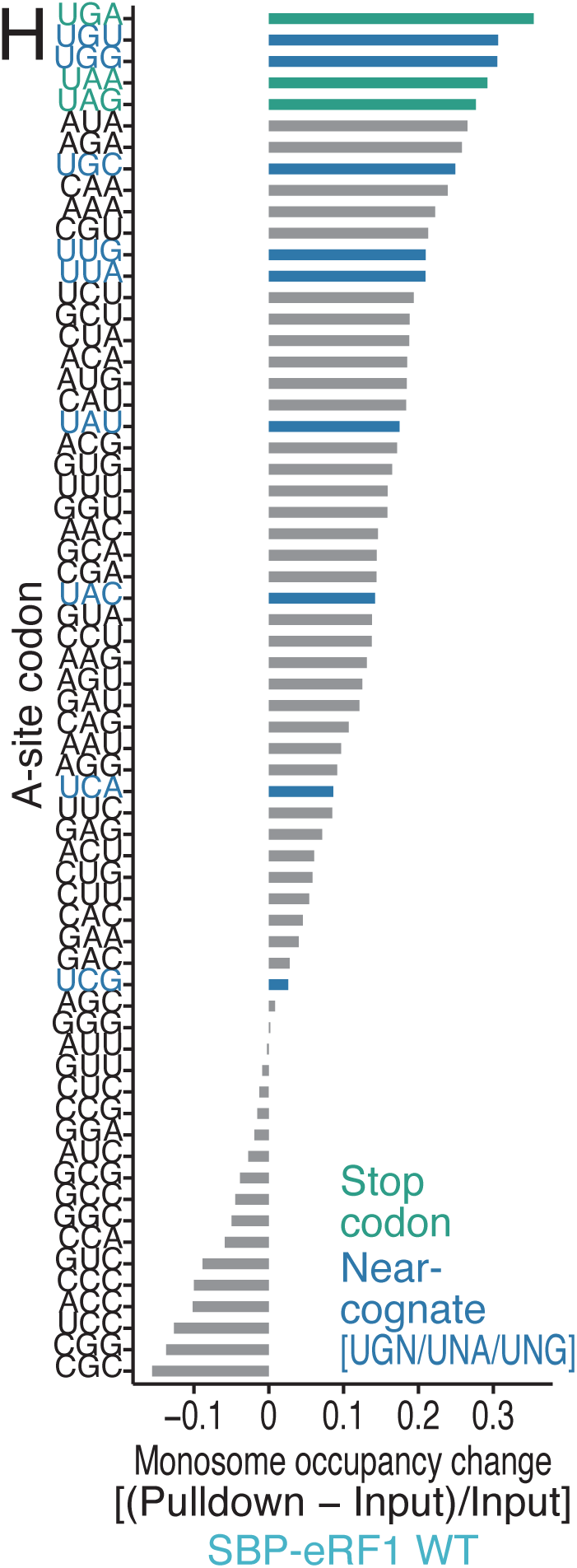
Characterization of eRF1-selective Monosome-Seq data, related to Figure 1. (A) Western blotting results for the indicated proteins. β-Actin was used as a loading control. (B) Distribution of monosome footprint lengths under the indicated conditions. (C) Heatmap of the Pearson’s correlation coefficients (*r*) of Monosome-Seq under the indicated conditions. The value of *r* is indicated by the color scale. (D) Fraction of the frame position for the 5′ end of the footprint (29 nt) under the indicated conditions. (E) Discrete Fourier transform of the 29-nt footprints whose 5′ ends mapped to the region from 150 nt upstream to 30 nt upstream of the stop codon. The signals of 30-nt periodicity (the size of the ribosome) may come from ribosome collision ^36^. (F) Plots of the 5′ ends of the monosome footprints around the stop codon (the first nucleotide of the stop codon was set to 0) under the indicated conditions. The color scale indicates the read abundance. RPM, reads per million mapped reads. (G) Comparison of the replicates for monosome occupancy differences between the eRF1 WT-bound fraction and the input. *r*, Pearson’s correlation coefficient; ρ, Spearman’s rank correlation coefficient. (H) Comparison of the replicates for monosome occupancy differences between the eRF1 WT-bound fraction and the input.

**Figure S2.**
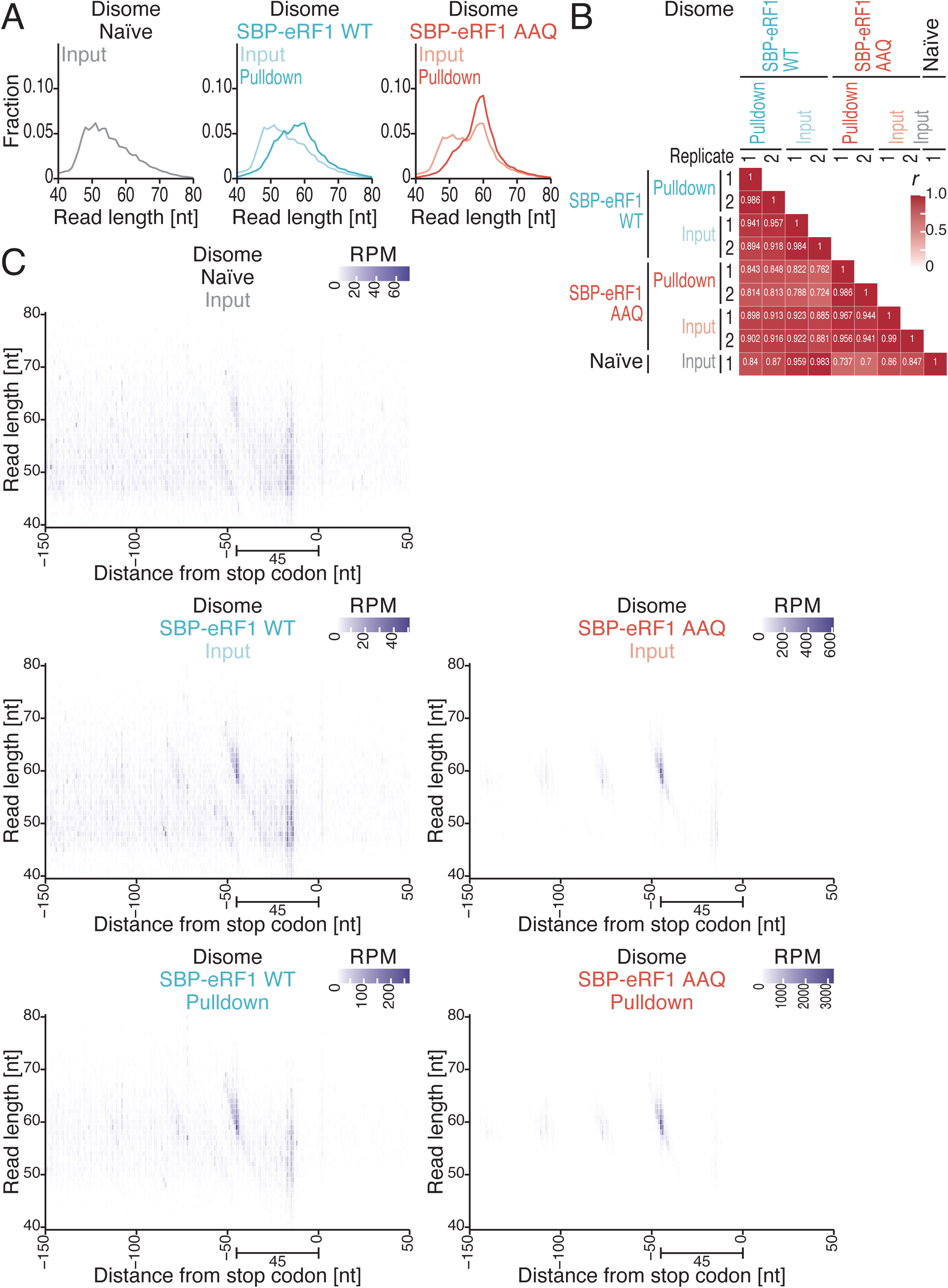
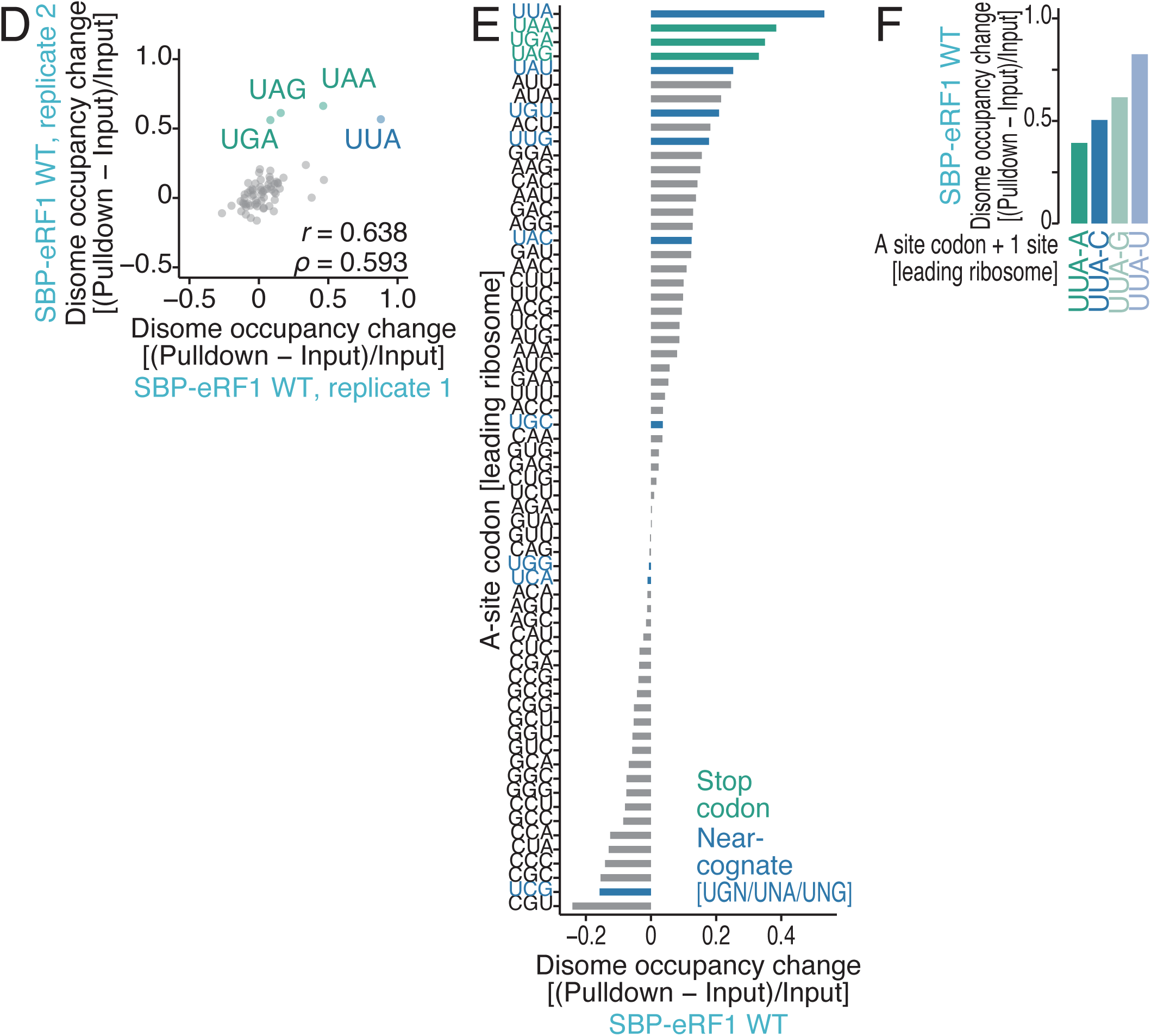
Characterization of eRF1-selective Disome-Seq data, related to Figure 1. (A) Distribution of disome footprint lengths under the indicated conditions. (B) Heatmap of the Pearson’s correlation coefficients (*r*) of Disome-Seq under the indicated conditions. The value of *r* is indicated by the color scale. (C) Plots of the 5′ ends of the disome footprints around the stop codon (the first nucleotide of the stop codon was set to 0) under the indicated conditions. The color scale indicates the read abundance. RPM, reads per million mapped reads. (D) Comparison of the replicates for disome occupancy differences between the eRF1 WT-bound fraction and the input. *r*, Pearson’s correlation coefficient; ρ, Spearman’s rank correlation coefficient. (E) Disome occupancy differences between the eRF1 WT-bound fraction and the input. (F) Disome occupancy differences between the eRF1 WT-bound fraction and the input for the indicated codon context.

**Figure S3.**
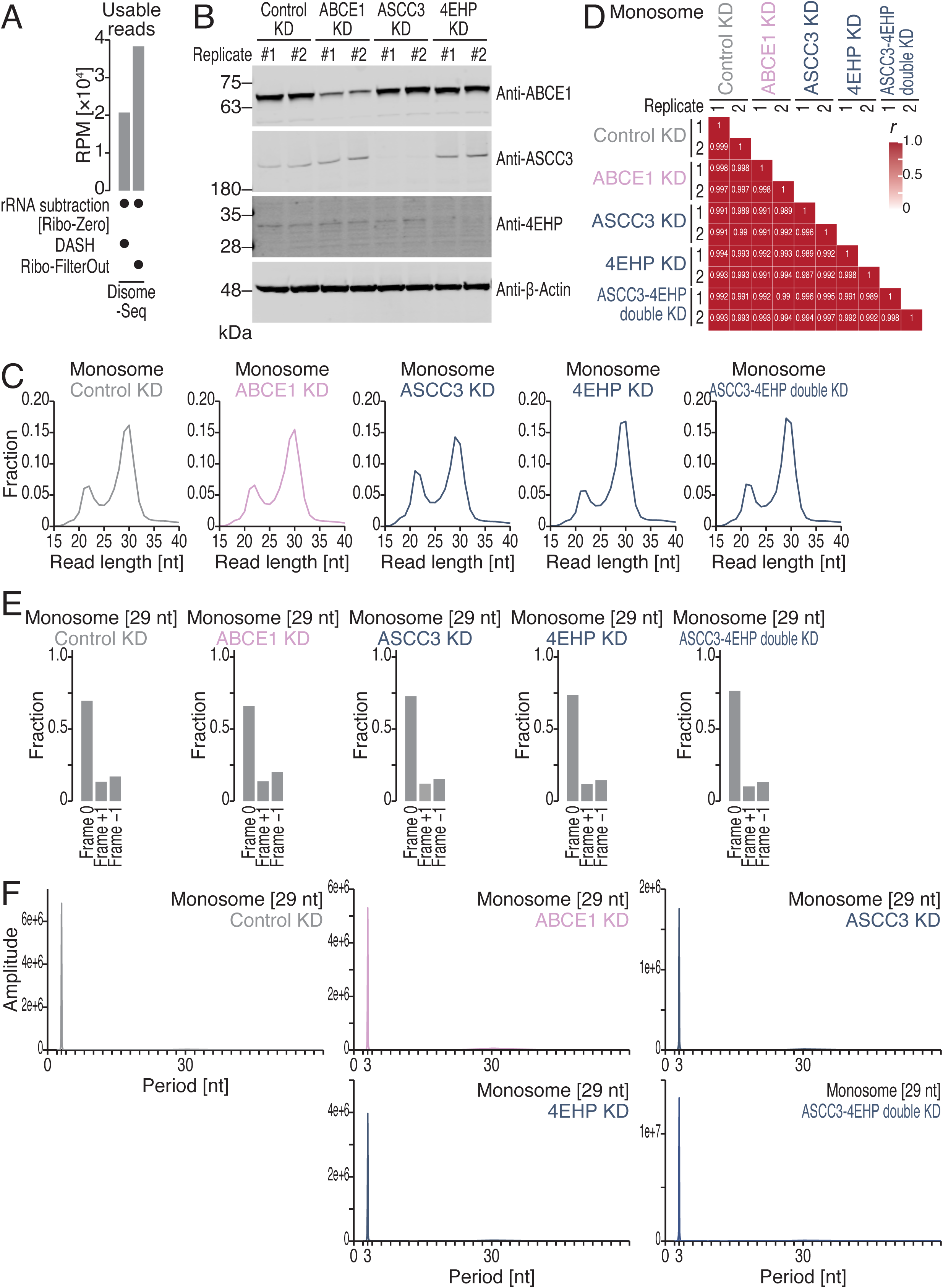
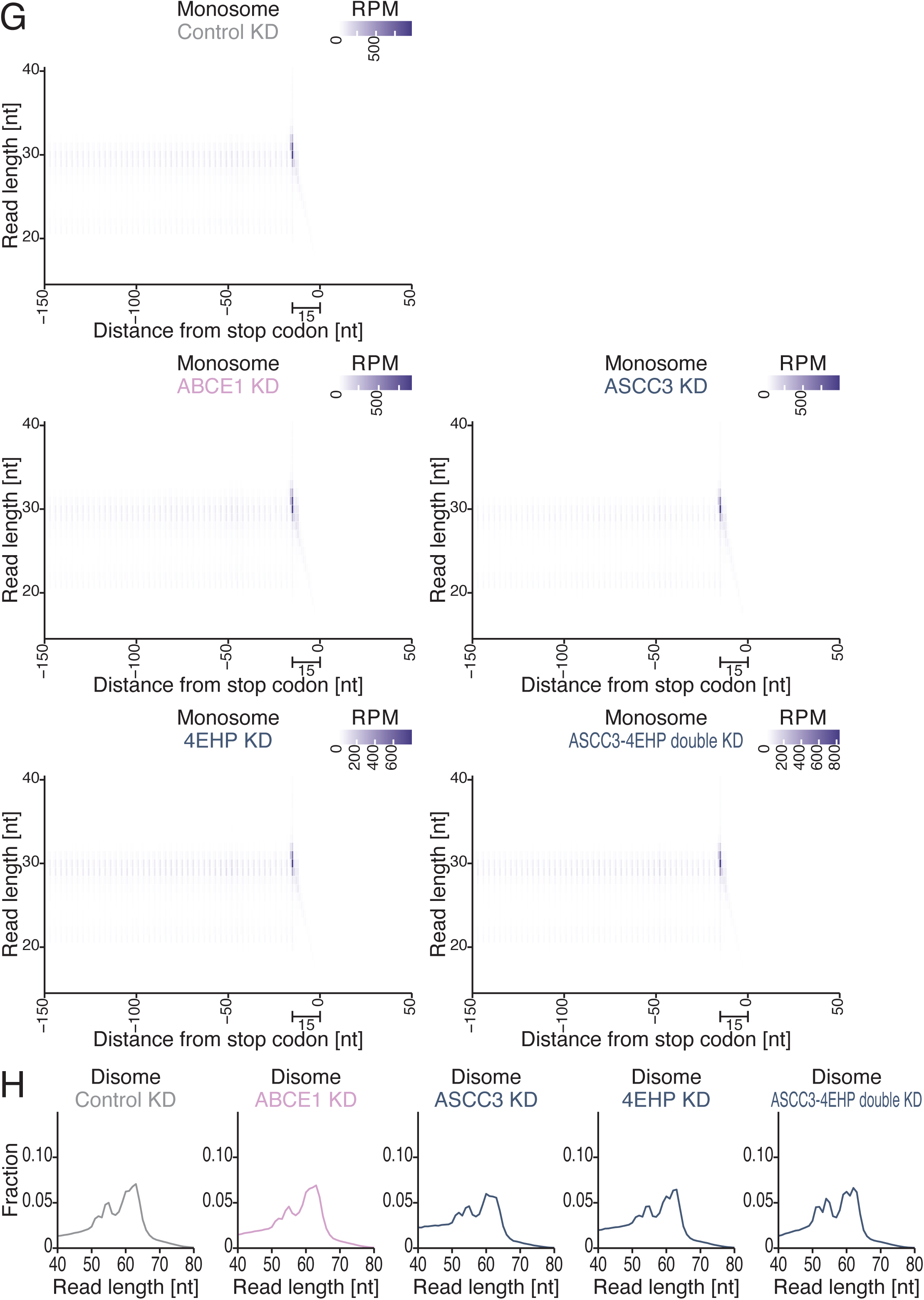
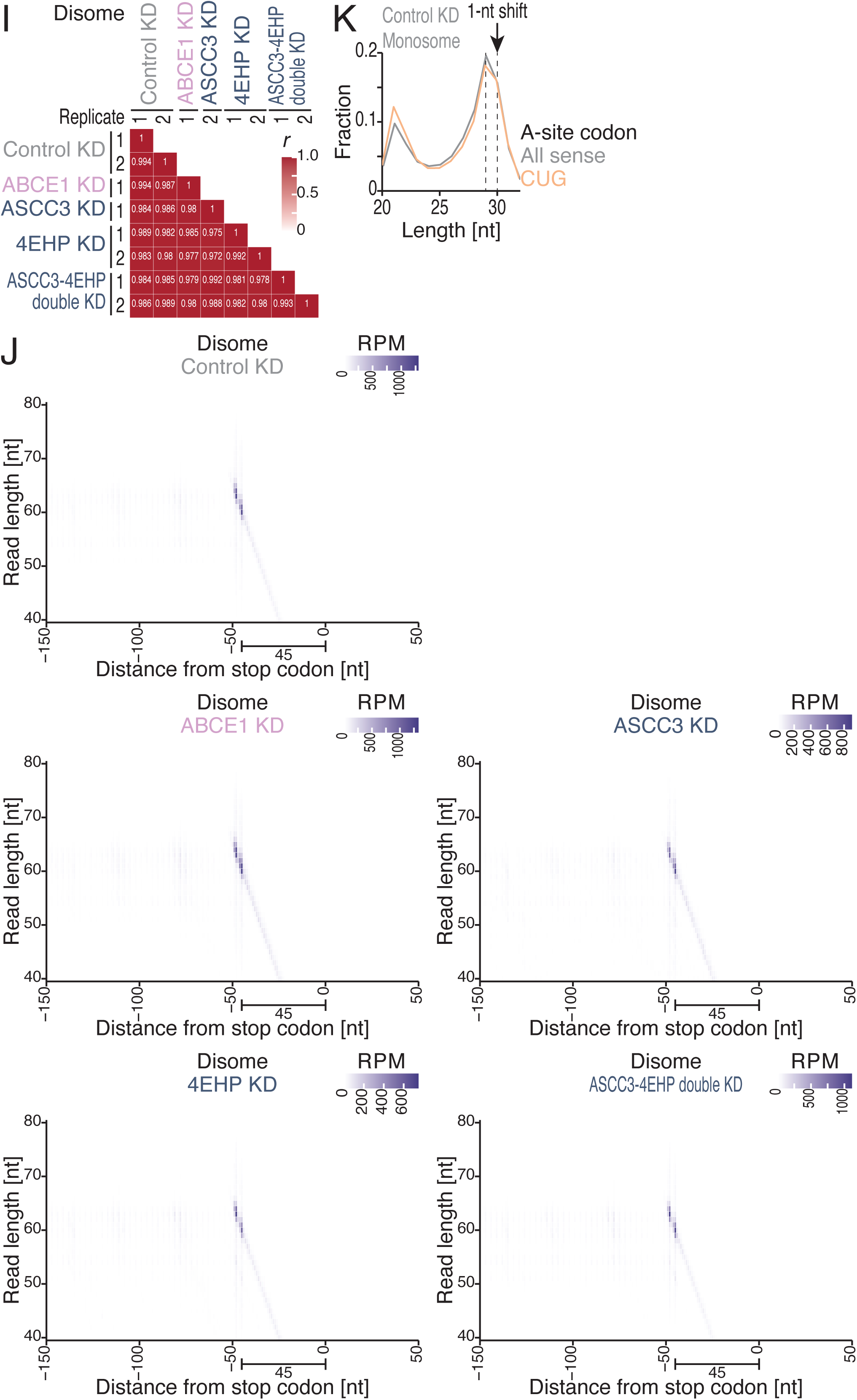
Characterization of Thor-Monosome-Seq and Thor-Disome-Seq data upon gene knockdown, related to Figures 1 and 3. (A) Fractions of usable reads (*i.e.*, genome-mapped reads, after deduplication and removal of noncoding RNA-mapped reads) for the indicated conditions for Disome-Seq library preparation. The DASH-treated data have previously been reported ^36^. (B) Western blotting results for the indicated proteins. β-Actin was used as a loading control. (C) Distribution of monosome footprint lengths under the indicated conditions. (D) Heatmap of the Pearson’s correlation coefficients (*r*) of Monosome-Seq under the indicated conditions. The value of *r* is indicated by the color scale. (E) Fraction of the frame position for the 5′ end of the footprint (29 nt) under the indicated conditions. (F) Discrete Fourier transform of the 29-nt footprints whose 5′ ends mapped to the region from 150 nt upstream to 30 nt upstream of the stop codon. (G) Plots of the 5′ ends of the monosome footprints around the stop codon (the first nucleotide of the stop codon was set to 0) under the indicated conditions. The color scale indicates the read abundance. RPM, reads per million mapped reads. (H) Distribution of disome footprint lengths under the indicated conditions. (I) Heatmap of the Pearson’s correlation coefficients (*r*) of Disome-Seq under the indicated conditions. The value of *r* is indicated by the color scale. (J) Plots of the 5′ ends of the disome footprints around the stop codon (the first nucleotide of the stop codon was set to 0) under the indicated conditions. The color scale indicates the read abundance. RPM, reads per million mapped reads. (K) Distribution of monosome footprint length at the indicated codon species in the A site.

**Figure S4.**
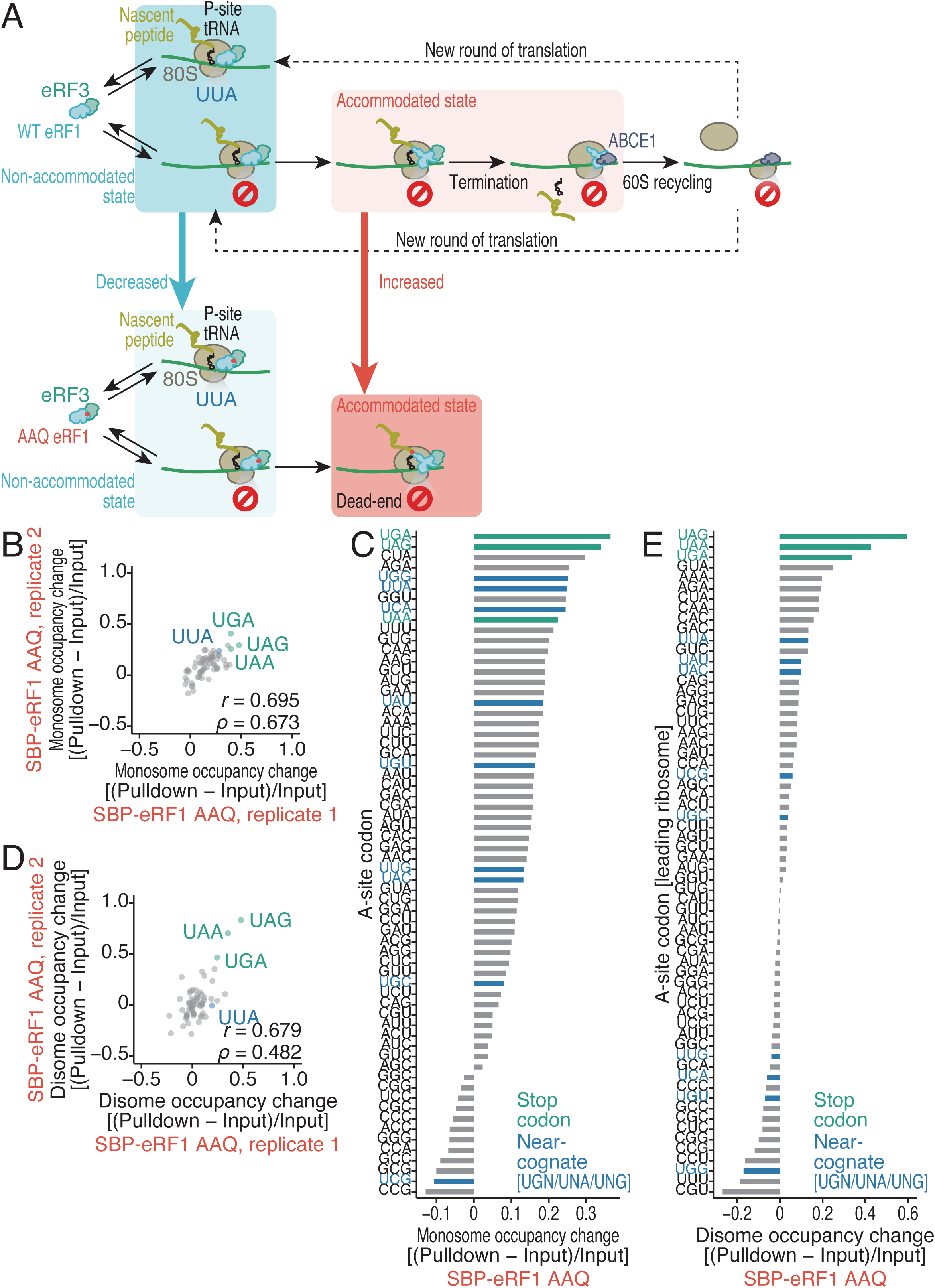
Effects of the eRF1 AAQ mutation on eRF1-selective Monosome-Seq and Disome-Seq data, related to Figure 1. (A) Schematic of the outcome of eRF1 (WT or AAQ mutant) association with the sense UUA codon and stop codon. (B and D) Comparison of the replicates for monosome (B) and disome (D) occupancy differences between the eRF1 AAQ-bound fraction and the input. *r*, Pearson’s correlation coefficient; ρ, Spearman’s rank correlation coefficient. (C and E) Monosome (C) and disome (E) occupancy differences between the eRF1 AAQ-bound fraction and the input.

**Figure S5.**
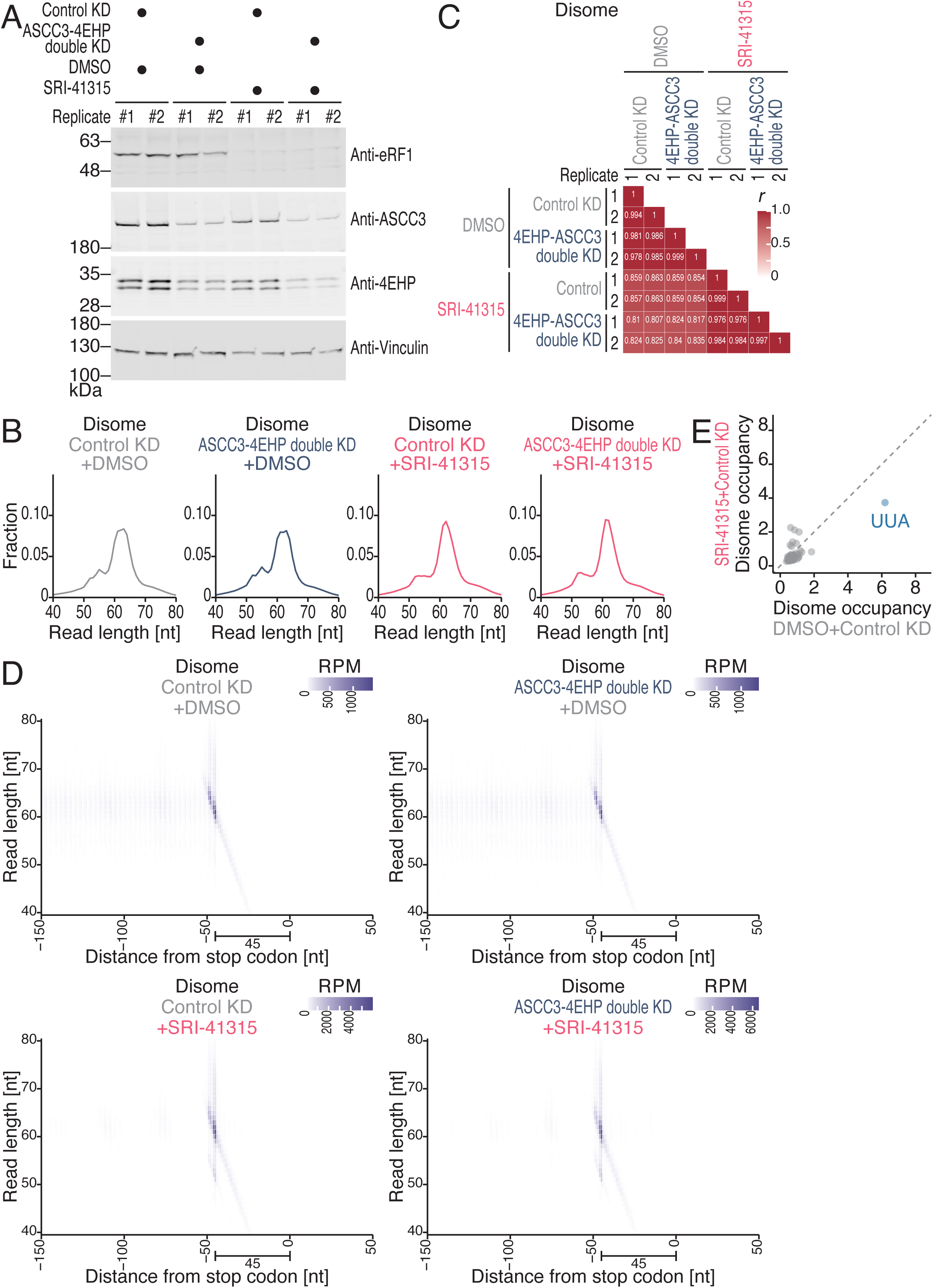
Characterization of Thor-Monosome-Seq and Thor-Disome-Seq data upon eRF1 depletion by the degrader compound, related to Figures 2 and 3. (A) Western blotting results for the indicated proteins. Vinculin (E) was used as a loading control. (B) Distribution of disome footprint lengths under the indicated conditions. (C) Heatmap of the Pearson’s correlation coefficients (*r*) of Disome-Seq under the indicated conditions. The value of *r* is indicated by the color scale. (D) Plots of the 5′ ends of the disome footprints around the stop codon (the first nucleotide of the stop codon was set to 0) under the indicated conditions. The color scale indicates the read abundance. RPM, reads per million mapped reads. (E) Comparison of disome occupancy on A-site codons (for the leading ribosomes) for the indicated conditions. Data for sense codons are shown.

**Figure S6.**
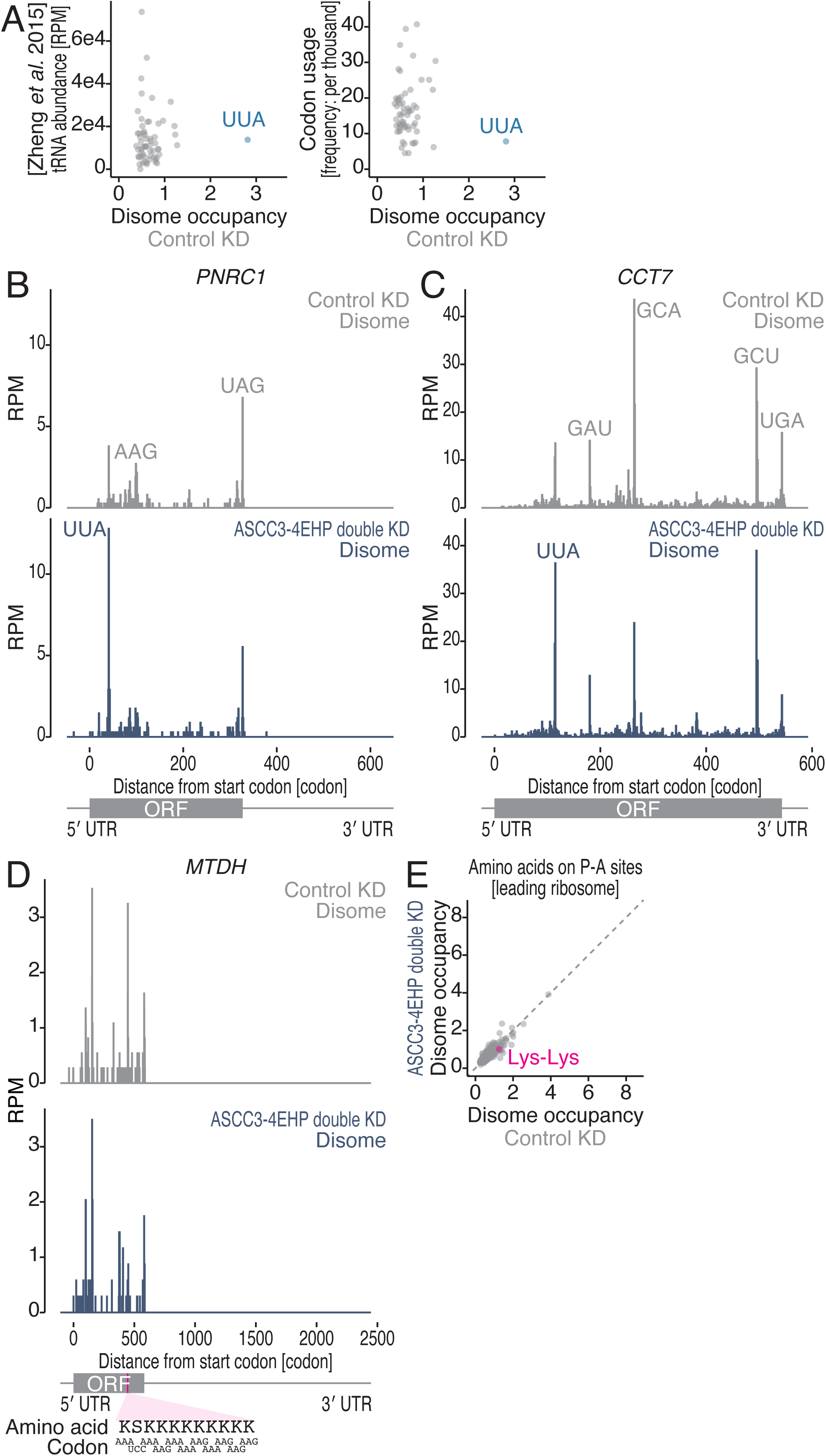
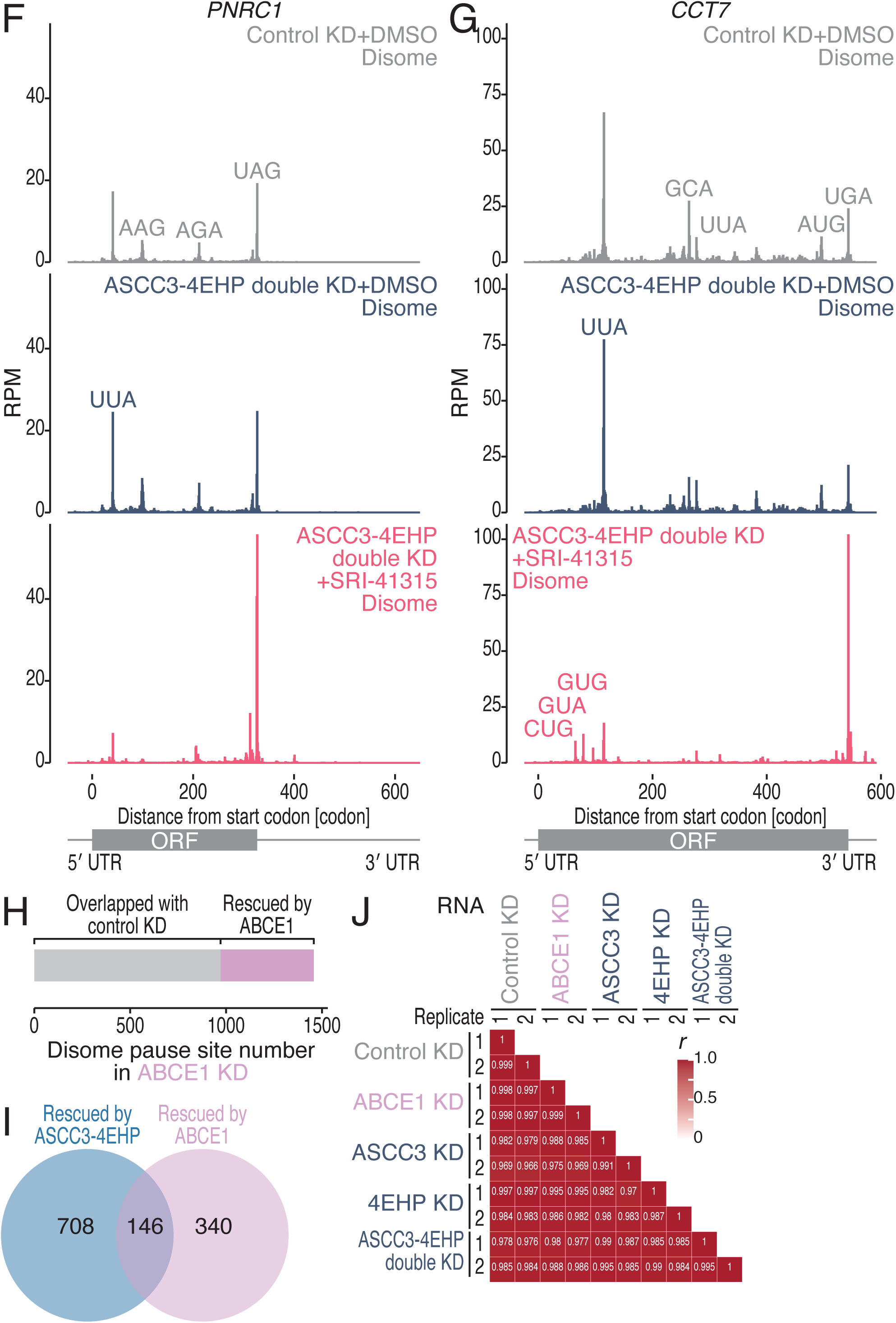
Characterization of ribosome collisions found in gene depletion, related to Figures 3 and 4. (A) Comparison of disome occupancy at A-site codons (for the leading ribosomes) with its cognate tRNA abundance measured by tRNA-Seq ^79^ (left) and the codon usage in the human genome (right). (B-D and F-G) Distribution of disome footprints along the indicated genes in the indicated conditions. The A-site position of the leading ribosome is depicted. The Lys-rich region is highlighted. (E) Comparison of disome occupancy on P- and A-site codon combinations (for disomes, A-site codons for the leading ribosomes) for the indicated conditions. The data were aggregated into amino acid pairs. (H) A breakdown of disome pause sites, illustrating the proportions of the indicated categories. (I) Venn diagram of disome pause sites found in the indicated conditions. (J) Heatmap of the Pearson’s correlation coefficients (*r*) of RNA-Seq under the indicated conditions. The value of *r* is indicated by the color scale.

**Table S1. Disome pause sites are defined in control siRNA-treated cells, related to Figure 3.**

mRNA name, codon position (the start codon is set as 0), raw read count, reads per million mapped reads (RPM), disome occupancy, ORF length (codon), gene name, pause score rank, ORF sequences in codon and amino acid, and E/P/A-site codon and amino acid sequences are listed.

**Table S2. Disome pause sites are defined in ABCE1 knockdown cells, related to Figure 3.**

mRNA name, codon position (the start codon is set as 0), raw read count, reads per million mapped reads (RPM), disome occupancy, ORF length (codon), gene name, pause score rank, ORF sequences in codon and amino acid, and E/P/A-site codon and amino acid sequences are listed.

**Table S3. Disome pause sites are defined in 4EHP-ASCC3 double knockdown cells, related to Figure 3.**

mRNA name, codon position (the start codon is set as 0), raw read count, reads per million mapped reads (RPM), disome occupancy, ORF length (codon), gene name, pause score rank, ORF sequences in codon and amino acid, and E/P/A-site codon and amino acid sequences are listed.

**Table S4. Disome pause sites are defined in 4EHP knockdown cells, related to Figure 3.**

mRNA name, codon position (the start codon is set as 0), raw read count, reads per million mapped reads (RPM), disome occupancy, ORF length (codon), gene name, pause score rank, ORF sequences in codon and amino acid, and E/P/A-site codon and amino acid sequences are listed.

**Table S5. Disome pause sites are defined in ASCC3 knockdown cells.**

mRNA name, codon position (the start codon is set as 0), raw read count, reads per million mapped reads (RPM), disome occupancy, ORF length (codon), gene name, pause score rank, ORF sequences in codon and amino acid, and E/P/A-site codon and amino acid sequences are listed.

